# A flexible intracortical brain-computer interface for typing using finger movements

**DOI:** 10.1101/2024.04.22.590630

**Authors:** Nishal P. Shah, Matthew S. Willsey, Nick Hahn, Foram Kamdar, Donald T. Avansino, Chaofei Fan, Leigh R. Hochberg, Francis R. Willett, Jaimie M. Henderson

**Affiliations:** Department of Neurosurgery, Stanford University; Department of Neurosurgery, The University of Texas at Austin, Austin, TX, USA + This work was primarily conducted at Stanford University.; Howard Hughes Medical Institute at Stanford University, Stanford, CA, USA; VA RR&D Center for Neurorestoration and Neurotechnology, Rehabilitation R&D Service, Providence VA Medical Center, Providence, RI, USA; School of Engineering, Brown University, Providence, RI, USA; Robert J. and Nancy D. Carney Institute for Brain Science, Brown University, Providence, RI, USA; Center for Neurotechnology and Neurorecovery, Department of Neurology, Massachusetts General Hospital, Harvard Medical School, Boston, MA, USA; Wu Tsai Neurosciences Institute, Stanford University, Stanford, CA, USA; Bio-X Institute, Stanford University, Stanford, CA, USA

## Abstract

Keyboard typing with finger movements is a versatile digital interface for users with diverse skills, needs, and preferences. Currently, such an interface does not exist for people with paralysis. We developed an intracortical brain-computer interface (BCI) for typing with attempted flexion/extension movements of three finger groups on the right hand, or both hands, and demonstrated its flexibility in two dominant typing paradigms. The first paradigm is “point-and-click” typing, where a BCI user selects one key at a time using continuous real-time control, allowing selection of arbitrary sequences of symbols. During cued character selection with this paradigm, a human research participant with paralysis achieved 30-40 selections per minute with nearly 90% accuracy. The second paradigm is “keystroke” typing, where the BCI user selects each character by a discrete movement without real-time feedback, often giving a faster speed for natural language sentences. With 90 cued characters per minute, decoding attempted finger movements and correcting errors using a language model resulted in more than 90% accuracy. Notably, both paradigms matched the state-of-the-art for BCI performance and enabled further flexibility by the simultaneous selection of multiple characters as well as efficient decoder estimation across paradigms. Overall, the high-performance interface is a step towards the wider accessibility of BCI technology by addressing unmet user needs for flexibility.

## Introduction

Brain-computer interfaces (BCIs) show great promise for restoring communication in people with paralysis (Mugler et al. 2014; Bacher et al. 2015; Herff et al. 2015; Gilja et al. 2015; Jarosiewicz et al. 2015; Pandarinath et al. 2017; Nuyujukian et al. 2018; Herff et al. 2019; Anumanchipalli, Chartier, and Chang 2019; Moses et al. 2021; Willett et al. 2021, 2023; Metzger et al. 2023; Card et al. 2023). User surveys for assistive devices have highlighted the need for multiple interface options that users can choose from based on their personal needs and capabilities (Scherer et al. 2005; Blabe et al. 2015; Fried-Oken, Mooney, and Peters 2015; Pitt and Brumberg 2018), encouraging new, previously unexplored directions for BCI research for communication. We introduce a typing BCI that can be used flexibly, allowing multiple high-performance options for communication.

The flexibility of finger movements in able-bodied people (Ingram et al. 2008; Xu, Mawase, and Schieber 2024) has contributed to the ubiquity of keyboards for typing on computers, driven in part by the different ways in which people can use them. For example, while trained touch-typists can achieve speeds of up to 200 words per minute, novice typists can still use the full capability of a QWERTY keyboard with one finger. Motivated by the recent demonstrations of decoding dexterous finger movements from intracortical recordings in monkeys (Nason et al. 2021; Willsey et al. 2022; Costello et al. 2023) and human participants (Wodlinger et al. 2015; Jorge et al. 2020; Vargas-Irwin et al. 2022; Guan et al. 2023; Shah et al. 2023; Willsey et al. 2024), we designed an intracortical BCI keyboard for typing based on flexion/extension movements of fingers. Focusing on finger movements simplifies the decoder algorithm compared with a QWERTY keyboard design, which would require decoding wrist movements in addition to finger movements.

Two distinct paradigms have emerged for BCI communication. The first paradigm, which we call “point-and-select” (Gilja et al. 2012; Bacher et al. 2015; Gilja et al. 2015; Kao et al. 2017; Pandarinath et al. 2017), involves real-time continuous control of a cursor with visual feedback (i.e., in closed-loop) to move it over a key before selecting it (e.g., with a discrete ‘click’ movement). By accurately selecting one character at a time, this provides a general-purpose typing paradigm that can be adapted to different languages, or even non-linguistic communication such as math equations, emojis, or musical notes without retraining the decoding algorithm.

The second paradigm involves decoding a rapid sequence of discrete movements with no immediate visual feedback (i.e., in open-loop), such as handwriting (Willett et al. 2021) or speech (Willett et al. 2023; Metzger et al. 2023). This enables high-speed handwriting and speech BCIs but relies on accurate movement decoding or error correction using the statistics of natural language.

We developed an intracortical BCI keyboard interface for selecting characters using flexion/extension finger movements and tested it in a BrainGate2 clinical trial participant (‘T5’) with two 96-channel silicon microelectrode arrays placed in the hand knob area (Yousry et al. 1997) of the left precentral gyrus (Fig. 1A). This interface simultaneously enabled high typing performance for both paradigms, simultaneous selection of multiple characters, bimanual finger movements from the unilateral implant and efficient decoder parameter estimation by exploiting the shared finger movements across tasks. This work presents a step towards personalized BCIs that can be adapted to individuals with different preferences, needs, and capabilities.

### An optimized BCI keyboard layout for typing with multiple fingers

**Fig. 1.**
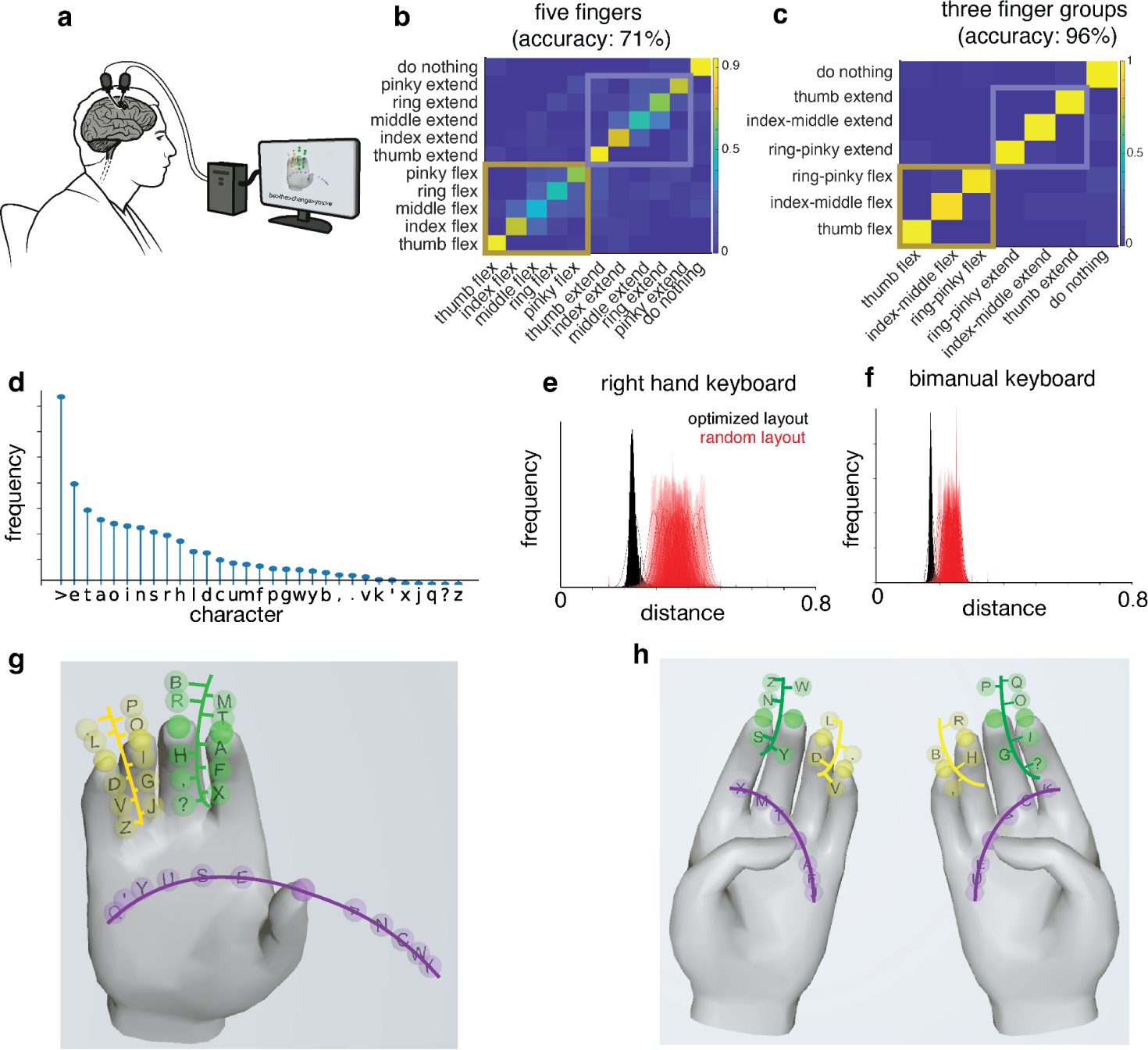
BCI keyboard design for typing with finger movements. (A) Neural activity was recorded using two ‘Utah’ arrays placed in the hand knob area of the left precentral gyrus as the research participant T5 attempted cued finger movements. (B) Confusion matrix (rows indicate cued movements and columns indicate decoded movements) for discriminating between single finger flexion/extension movements using a naive Bayes classifier. Trials were open-loop, with the participant instructed to attempt to follow an animated hand. Following the instructed delay paradigm, each trial had 1s preparatory time, 1s movement time, and 0.5s hold time. (C) Confusion matrix for a reduced set of movements with three finger groups corresponding to the thumb, index-middle tied together, and ring-little tied together. Each trial had 1s preparatory time, 0.5s movement time and 1s hold time. Reducing the movements to three degrees of freedom improved classification accuracy. (D) The relative frequency of the 31 symbols (26 English letters, space indicated by ‘>’, and four symbols) used in this paper was estimated from 57,340 sentences of the Brown corpus (Francis and Kucera 1979). (E, F) Distribution of mean distance traveled per character for sentences from the Brown corpus comparing the optimized keyboard layout (black) to random layouts (red) for the right hand (E) and bimanual keyboard (F). Distance normalized such that going from complete flexion to extension is equal to 1. (G) Right-hand keyboard layout with three degrees of freedom. Colors indicate finger groups. Index-middle fingers (green) are constrained to move together. Similarly, ring-little fingers (yellow) are constrained to move together. Keys are laid out along the flexion-extension axis for each finger group. Keys along the two fingers belonging to a finger group are staggered to enable the selection of a unique key with constrained movements. (H) Bimanual keyboard layout with six degrees of freedom and three finger groups per hand.

Ideally, a typing keyboard would be based on isolated movements of all 5 individual fingers. While the attempted movements of single fingers can be decoded from neural recordings in T5, (71% accuracy, Fig. 1B), constraining nearby fingers to move together (thumb, index-middle tied together, and ring-little tied together) resulted in greater accuracy (95%, Figure 1C). Next, we designed a keyboard for selecting characters using the movement of three finger groups.

To construct our keyboard, we considered 26 letters, a symbol for space (>), and punctuation (question mark, comma, period, and apostrophe), resulting in a total of 31 symbols (Willett et al. 2021). Keys were arranged along the flexion/extension direction of each finger group and roughly equal numbers of keys were assigned to each of the six directions on the right hand (three finger groups x 2 movements per finger group). More frequent symbols in the English language were assigned to keys at a smaller distance from rest (Fig. 1D). For keys that were equidistant from rest but on different fingers, the symbols were first assigned to the thumb, followed by index-middle and ring-little finger groups. As a result, high-frequency symbols such as “E”, “T”, “A”, and space (“>”) are only a small movement away from the rest position of the corresponding finger.

Under the assumption of constant velocity throughout a trial and independent control of individual finger groups, the typing rate is proportional to the average distance traveled per character. For a particular keyboard layout, the average distance per character was computed for all characters in a sentence, and the distribution across all sentences was evaluated using the Brown corpus (Francis and Kucera 1979). The distribution for the optimized keyboard (black, Fig. 1E) was substantially smaller than the distribution for different random letter layouts (red, Fig. 1E). This optimization was repeated for the bimanual keyboard with the keys laid out across 12 directions on both the hands, resulting in an even smaller average distance traveled (Fig. 1F) and fewer keys per finger group (Fig. 1H).

### “Point-and-click” character selection with closed-loop finger movements

Point-and-click typing consists of moving a pointer to a target key using closed-loop control and then selecting the character with a discrete click signal. In our case, the pointer corresponds to the position of a virtual fingertip and clicking corresponds to an elbow or ankle flexion movement, depending on the task (see below for details).

Point-and-click typing involves the use of two distinct decoders: a decoder for guiding virtual finger movements and a click decoder for initiating discrete clicks. The click decoder, based on logistic regression, was trained using open-loop trials performed without visual feedback. During these trials, participant T5 performed finger movements followed by the click movement. Concurrently, a finger movement decoder was trained on closed-loop point-and-hold trials. In these trials, T5’s task was to maneuver a finger over a designated key (’point’) and then maintain its position over the key for two seconds (’hold’).

The velocity decoder was developed using a recipe that addresses some of the fundamental challenges behind high degree of freedom motor decoding. First, as the distributions of neural activity between open-loop (without feedback) and closed-loop (with real-time visual feedback) differ significantly (Fig. 2A, left), we collected training data in closed-loop using a simple logistic regression-based decoder which was updated in real-time (Fig. 2A, right). Second, as the finger movements are represented non-linearly in the motor cortical neural activity (Willsey et al. 2022; Shah et al. 2023), we used a multi-layered feedforward neural network to decode instantaneous finger velocities from the recorded neural activity (architecture shown in Fig. 2B, adapted from (Willsey et al. 2024)). While real-time continuous control was also achievable by directly decoding finger positions, velocity decoding was faster (Fig. S1, Videos S1, S2). Third, as neural network models with a large number of parameters (like those considered here) require a large training dataset, in an effort to reduce in-session calibration times, the network was pretrained using the training data from previous sessions and fine-tuned using training data from the target session (Fig. 2C). Fourth, as the integration of noisy velocity predictions for a non-target finger results in position drift and requires corrective movements resulting in longer trials, soft-thresholding (which makes all velocity predictions less than a certain value exactly zero) provides improved ability to keep the non-target fingers at rest (Fig. 2D, Fig. S2).

**Fig. 2.**
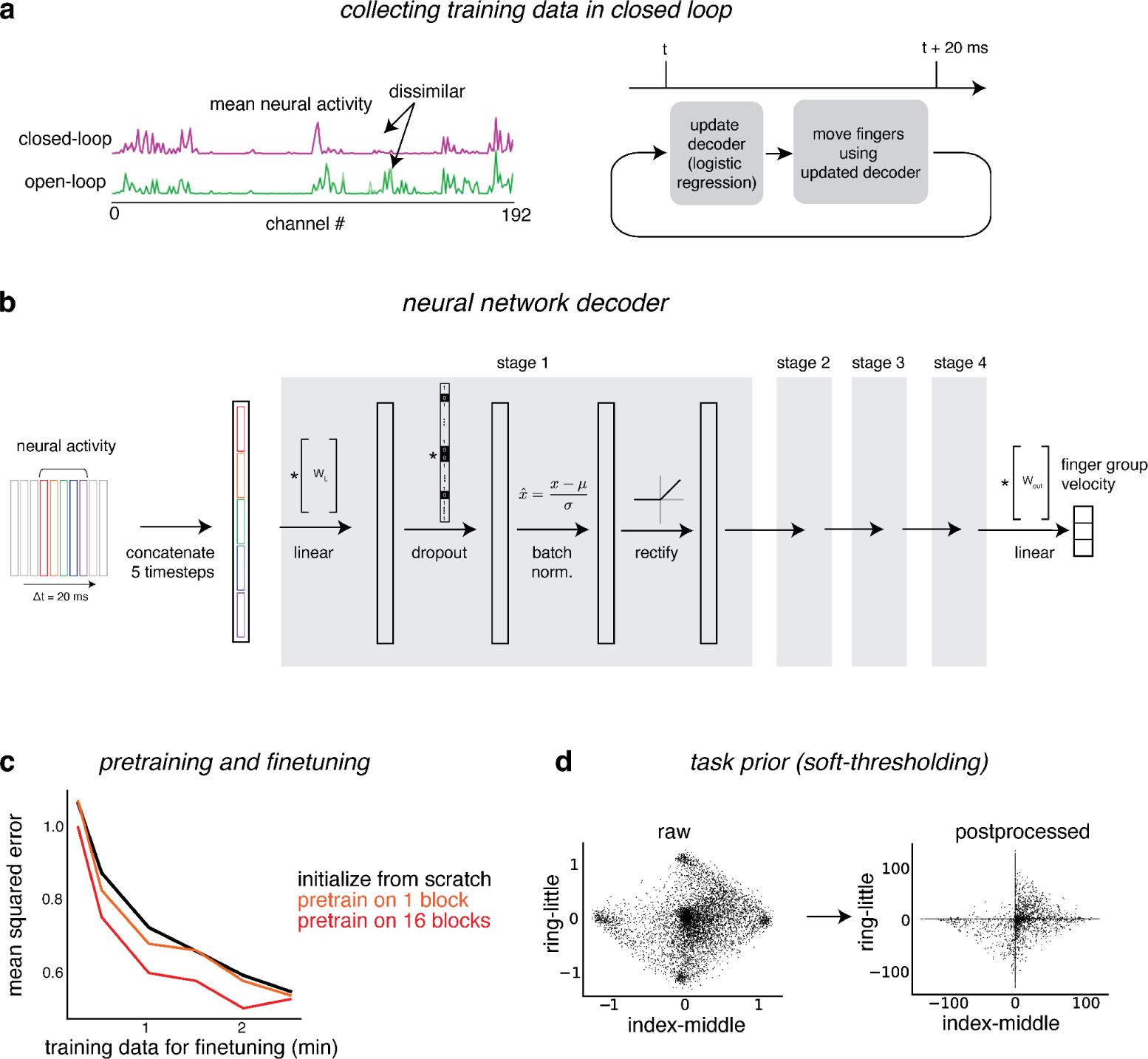
A streamlined recipe for neural network decoding for movements with multiple degrees of freedom. (A) (Left) Differences in the neural activity between open-loop (without visual feedback) and closed-loop (with real-time visual feedback) tasks. Lines indicate the mean number of threshold crossings across channels for a single block. Channels with a large difference in activity between open-loop and closed-loop blocks are highlighted with the black arrow. (Right) Training data collection using point-and-hover task, involving closed-loop control to point a finger over a key and hovering over a key for 2 seconds to select it. At each time step, a logistic regression-based decoder was updated using new data and immediately used to control finger movements. (B) Decoder architecture. Neural activity (concatenation of spike power and binned threshold crossings) from the previous five time steps (20ms bins) predicts finger velocities after passing through multiple stages of a linear layer, dropout, batch normalization, and rectified non-linearity. (C) Pretraining the decoding algorithm using data from the previous session improved decoder performance. Prediction mean-squared error on 5.4 to 9.4 min of held-out data (y-axis) with the amount of data used for training (x-axis). The decoder is either trained from scratch (i.e. random initialization) using data from the target session (black) or pre-trained using blocks from other sessions and then fine-tuned using data from the target session (other lines). Pretraining using either 1 previous session (orange) or 16 previous sessions (red). Performance was averaged over 5 partitions of data, with one randomly selected block used for testing, and either one or 16 other blocks were used for pretraining. (D) Predicted velocities for the index-middle and ring-little groups (each dot is a sample). Raw velocities indicated on left and post-processed (with gain, smoothness, and soft-thresholding f(x) = max(x-t, 0) - max(-x-t, 0), with threshold t) velocities indicated on right. More points are along the x-axis or the y-axis with soft-thresholding, as required by the task where only one finger group should be moving at a time.

Typing performance was evaluated using a cued character selection task, where the target characters were visually cued for every trial. Target characters for successive trials correspond to English sentences from (Willett et al. 2021). The task is designed to estimate an upper bound for ‘free typing speed’ by removing the requirement for the user to locate the desired key (in addition to controlling finger movement). True free typing speed would thus necessarily be slower than this upper bound due to the increased cognitive load of deciding on responses and locating each symbol on the keyboard.

When pointing with the right-hand keyboard and clicking with the left elbow, T5 achieved 31.7 (SD: 2.4) correct characters per minute (CCPM) and 89.1% (SD: 3.2%) success on average (Fig. 3D, E, F, Fig. S3, Video S3, See Table S1 for all blocks). Time per trial ranged between 1.19s for the nearest keys (comprising 51% of the targets) and 2.67s for the farthest keys (3% of the targets), resulting in an average trial time of 1.49s (Fig. 3F). To identify avenues for further improvement in speed, the total trial duration was decomposed into (1) the time taken to first reach the target and (2) selection time, which includes the time spent ‘orbiting’ the target and completing a click. While the time to first reach the target increased with distance as expected, a constant amount of time (∼0.5s) was spent orbiting and selecting the target key for each trial (Fig. 3F). Motivated by this observation, we explored two avenues for increasing the point-and-click typing speed.

**Fig. 3.**
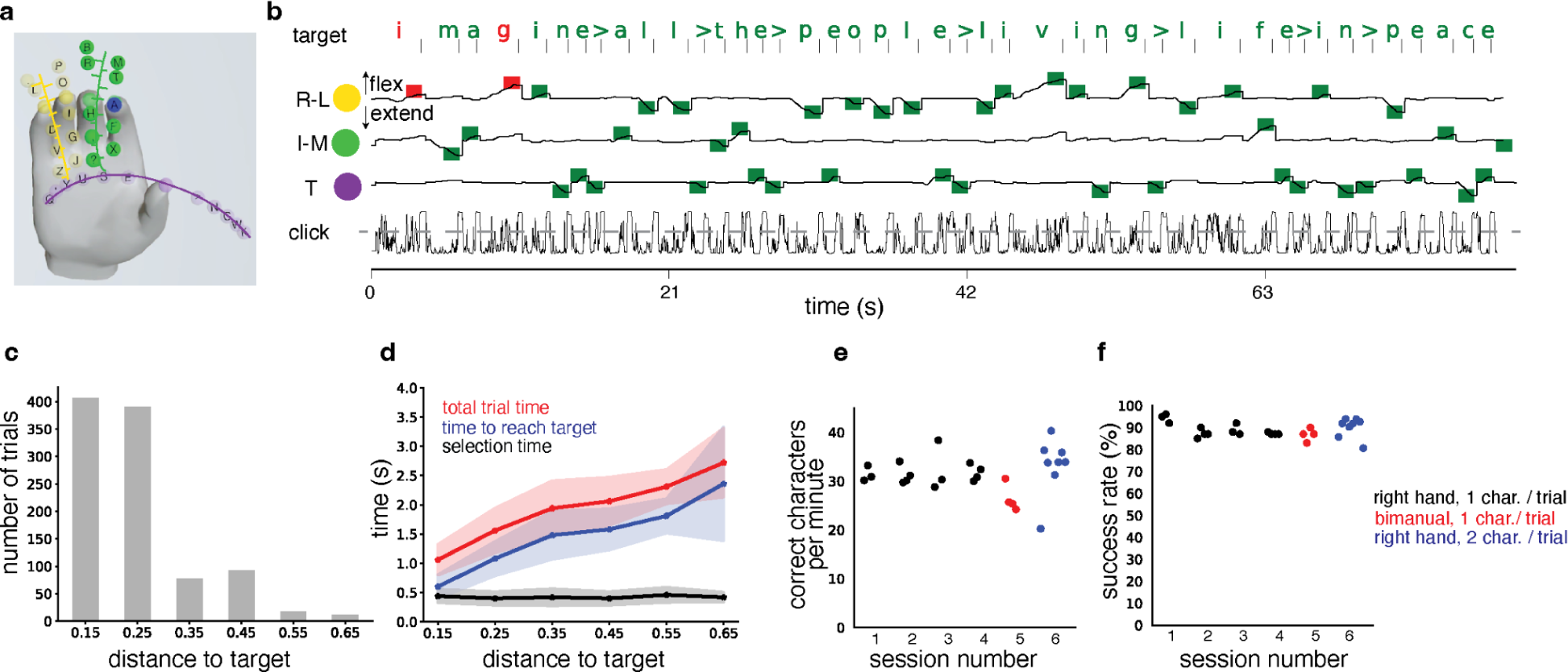
Point-and-click character selection on right-hand and bimanual keyboards. (A, B) Finger position trajectories for an example block for typing on the right hand with three finger groups. Finger groups are moved to the target positions (squares) and selected with a click corresponding to flexing the left elbow for selecting the character (click probabilities indicated at bottom, click threshold indicated with dashed line). Finger groups are denoted by T (thumb), I-M (index-middle), and R-L (ring-little). Correctly (incorrectly) selected characters indicated green (red). (C) Distribution of target distances for selecting English sentences with right-hand keyboard and 1 character selection per trial. Data was combined across 14 blocks of 100 trials each, collected on four different session days. Note that the number of trials decreases with increasing distance (as intended by the keyboard design). (D) Completion times for trials in (C). Total trial times, time to first reach the target, and the selection time (measured as the difference between total trial time and first reaching the target) are separated. Note that the total trial time and time to target increases with the distance of the target key and selection time does not change with the distance of the target key. Individual trials are shown in Fig. S3. (E, F) Correct characters per minute (E) and success rate (F) for point-and-click typing with the three-finger groups on the right hand (black), six-finger groups on both hands (three-finger groups on each hand, red) and simultaneous selection of up to two characters with a single click on the right hand (blue). Dots indicate performance computed from a block of 100 trials. X-labels indicate sessions.

The first approach aimed to reduce the average distance traveled by using a keyboard developed for both hands, resulting in six finger groups (three on each hand). Compared to the right-hand keyboard, the bimanual keyboard had fewer characters per finger (Fig. 1H). When T5 attempted to click with the left hand while using the bimanual keyboard, the click times were high, due to a lag in switching between the attempted movement of the fingers on the left hand and the movement of the left elbow (self-reported by T5). In contrast, clicking with an attempted “gas pedal” movement on both ankles was faster. However, compared to the right-hand keyboard, this strategy did not result in an improved overall typing performance (CCPM: 26.5 with SD 2.41, Accuracy: 86.8% with SD 2.5%), presumably because of greater noise in decoding the ipsilateral movements and larger number of degrees of freedom (Fig. 3E, F, Video S4).

The second approach aimed to reduce the total time spent clicking by enabling the selection of two successive characters at once. Specifically, if successive characters in a sentence are on different finger groups, they can be simultaneously pointed to by corresponding fingers and then selected with a single click. This approach has recently shown high throughput in a center-out target acquisition task using virtual fingers (Willsey et al. 2024). For the right-hand keyboard with three finger groups, 63% of trials had two successive characters on different finger groups for typing English sentences. While this approach exploits the participant’s ability to attempt movements of multiple fingers simultaneously, a language model is required to identify the correct order of the two simultaneously selected characters. Using this method, T5 was able to select characters at 33.2 (SD 5.48) characters per minute, a modest improvement from single character selection per trial (Fig. 3E, F, Video S5). Notably, a language model from (Fan et al. 2023) was able to identify the correct order of characters resulting in an overall 9% character error rate. Although this method allowed for slightly faster typing, it would likely require extensive practice given the much higher cognitive load required to select two consecutive characters at once. Regardless, throughput in this range is consistent with previous reports of point-and-click typing ranging from 31-40 correct cpm (Pandarinath et al. 2017).

### “Keystroke” character selection with open-loop finger movements

The second paradigm we considered is “keystroke typing”, which allows the selection of a sequence of characters with discrete finger movements performed in an open-loop (without feedback). In this approach, the magnitude of finger movement is ignored and only flexion versus extension is decoded. Ideally, the number of symbols must be matched to the number of discrete movements, so that the information transmission is lossless if the movements can be perfectly decoded. However, this is not possible given the need to encode 31 symbols with six discrete movements for the right-hand keyboard (with three finger groups and two flexion/extension movements each), resulting in confusion between 4-6 characters per movement, or 12 discrete movements for the bimanual keyboard (with three finger groups of each hand), resulting in a confusion between 2-3 characters per movement.

Even though perfectly lossless communication is infeasible, one could leverage the fact that most finger movement sequences likely correspond to a single realistic English sentence. Assuming a perfect ability to decode finger movements, the capability of such an error correction method was determined using the following analysis (Fig. 4A, B, C, D). First, sentences were encoded into a sequence of finger movement directions. Then, uniform probability was assigned to all characters along a finger movement direction. Finally, a language model (Fan et al. 2023) used the sequence of probabilities to identify the most likely English sentence. This procedure resulted in a 5% error for the right-hand keyboard and less than 1.5% error for the bimanual keyboard. Since the error rate was lower, we evaluated the performance of keystroke typing using the bimanual keyboard in participant T5.

**Fig. 4:**
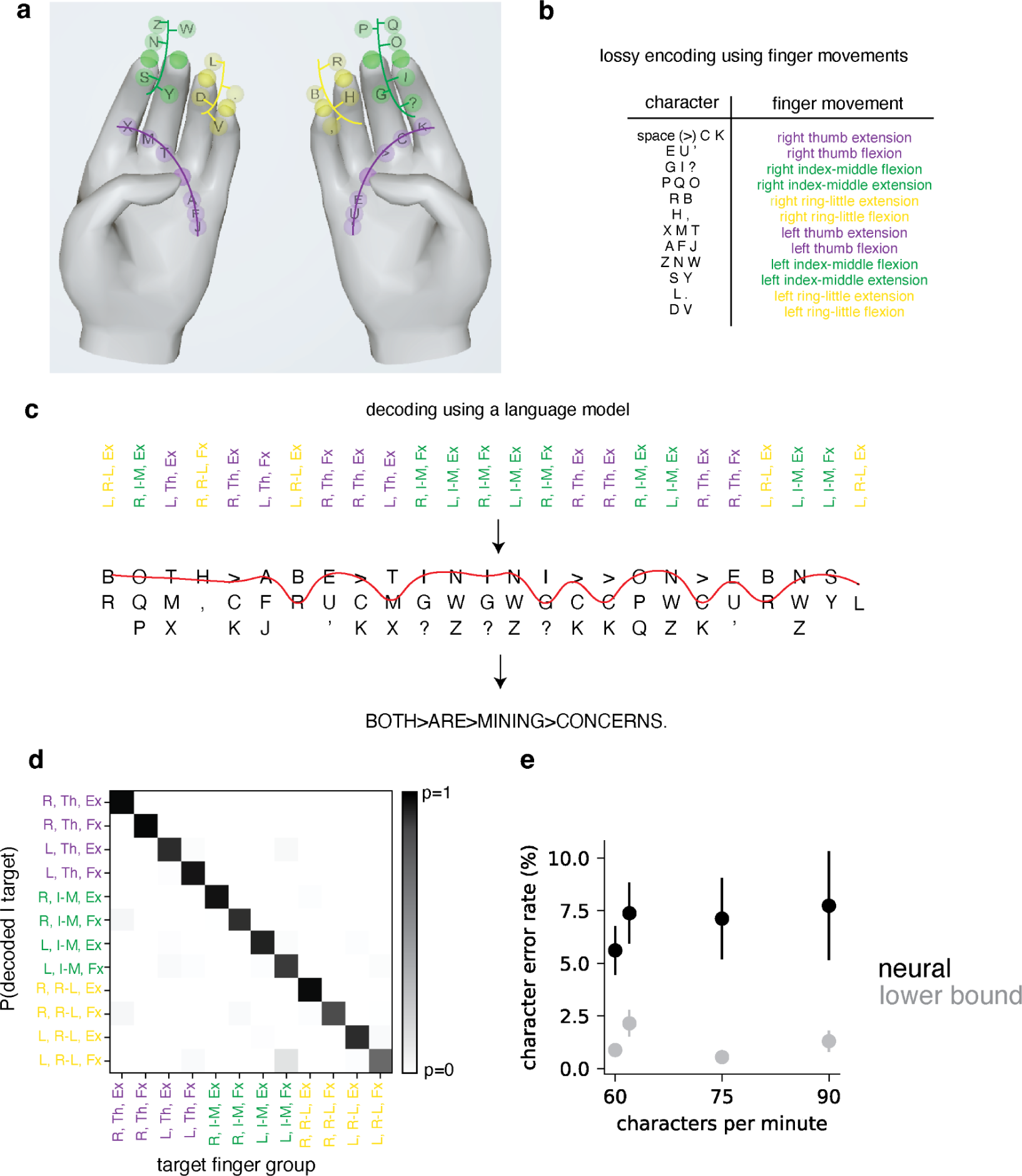
Character selection with rapid keystroke movements on the bimanual keyboard. (A) The bimanual keyboard reproduced from Fig. 1H. (B) Table shows the characters and corresponding finger group movement used for encoding them. Multiple characters map to the same finger movement, resulting in lossy encoding. (C) Decoding using a language model. First, finger movements are decoded from recorded neural activity (top row). Finger movements are indicated by a tuple of laterality (L/R), finger group (thumb: T, Index-Middle: I-M, Ring-Little: R-L), and movement (Extension: Ex, Flexion: Fx). Next, the subset of characters encoded by each finger movement is identified (second row, characters along a column sorted according to the decreasing probability in the English language). Third, a tri-gram language model (from (Fan et al. 2023)) is used to identify plausible English sentences from the decoded characters (the decoded sentence is indicated by the red line and the third row). Detailed formulation in Methods and Fig. S4. (D) Mean probability over fingers for different target fingers using a logistic regression-based decoder, averaged over 28 sentences presented at 1 second/character. (E) Character error rate after decoding finger movement and applying the language model (black) for sentences cued at different speeds (60, 75, or 90 characters per minute). Each dot represents a session. A lag of 0.25 sec was added to neural activity compared to cue onset for the fastest speed (0.67 sec/char). Lower bounds (gray) measured the best possible character error rates assuming that the target fingers are perfectly decoded. Specifically, all keys along a target finger direction are assigned a uniform probability and processed by a language model to identify the most likely sentence. Error bars indicate the standard deviation of the error rate, computed by bootstrapping the sentences.

Neural activity was recorded as T5 attempted a sequence of finger movements toward a target character, with a different cued character shown at regular intervals of 0.67 seconds, 0.8 seconds, or 1 second across tasks. For each movement, a logistic regression classifier predicted a probability distribution over finger movements. This probability distribution was then fed as input to the error correction procedure described above. For 1 second per character (two sessions), 0.8 second per character (one session) and 0.67 second per character (one session) speeds, the final character error rate was <10%, resulting in typing speeds of 60, 75, and 90 characters per minute respectively. These error rates were close to the best possible error rates assuming perfect finger decoding (Fig. 4E), suggesting a minimal impact of finger decoding error on final performance. Overall, the peak performance of “keystroke” typing is comparable to the high communication throughput seen in classifying handwritten characters at 90 characters per minute with error rates of <1% (Willett et al. 2021).

### Leveraging shared finger movements for efficient decoder estimation

The ability to use finger movements across two distinct typing paradigms (continuous and discrete) makes it possible to build a shared decoder across them. Foundation models, which have recently become popular in machine learning, are pretrained on a diverse collection of tasks and can be efficiently adapted (fine-tuned) on various downstream tasks to provide high accuracy (Bommasani et al. 2021). The possibility of such a foundation model for finger movements was tested in different ways.

First, we tested if a neural network decoder pre-trained on data from multiple sessions of point-and-hold typing with the right-hand keyboard improved decoding in a new session of the same task (Fig. 3D). While the performance of both a pre-trained decoder and a decoder trained from scratch increased with the amount of fine-tuning data from the target session, the performance of the pretrained decoder was consistently superior. Moreover, the performance gap increased with more data used for pretraining, suggesting that the neural network decoder identified useful features by combining data across multiple sessions.

Next, we tested if similar improvements were observed by combining data across related tasks. A logistic regression decoder already performed well for keystroke typing, so we only tested this approach for learning the neural network decoder for point-and-hold tasks. For the point-and-hold tasks with either the right hand or both hands, fine-tuning a decoder that was pre-trained on *all* the tasks (11 sessions of closed-loop right-hand point-and-hold, 7 sessions of closed-loop bimanual point-and-hold and 17 sessions of open-loop bimanual keystroke sequences) performed better compared to a decoder trained from scratch (i.e., randomly initialized parameters; Fig. 5A, B). For both open-loop and closed-loop tasks, the decoder training assumed that the participant was moving towards the cued keys at each point of time and the decoder architecture was modified to predict both the continuous-valued finger velocities and discrete finger groups (see Methods). Notably, combining data from the keystroke character selection task improved decoder performance compared to a decoder pre-trained only for the point-and-hold task. This suggests that combining data from multiple related tasks with shared underlying finger movements is beneficial.

**Fig. 5:**
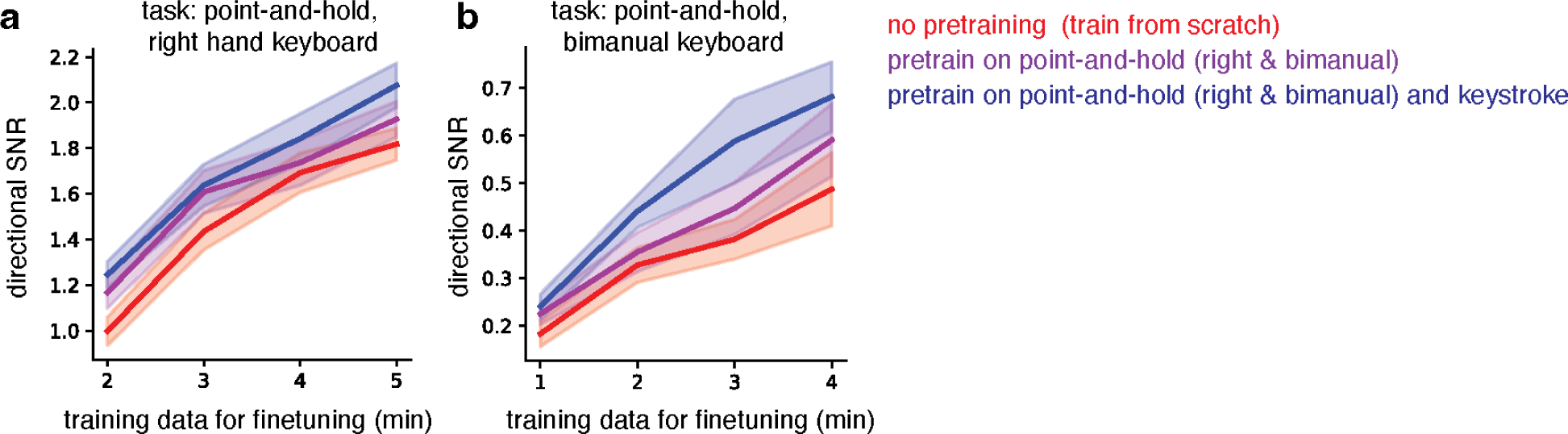
Leveraging shared finger movements across tasks enables efficient decoder estimation. (A) Performance (y-axis) of three decoders on the closed-loop point-and-hold task with the right hand with different amounts of training data (x-axis). Decoders were either pre-trained on other sessions and fine-tuned on the target session (purple and blue) or fine-tuned on the target session from scratch (red, random initialization of decoder parameters). Performance was evaluated using the directional signal-to-noise ratio (dSNR, (Willsey et al. 2024)) of predictions on testing data. The solid line indicates the average dSNR across ten blocks and transparent lines indicate the standard error across blocks. While the dSNR increased with the amount of fine-tuning data for all decoders, the pre-trained decoders (purple and blue) outperformed the decoder fine-tuned from scratch (red). Pretraining on the closed-loop point-and-hold as well as the open-loop keystroke typing task (blue) outperformed a decoder pre-trained only on the closed-loop point-and-hold task (purple). See the methods section for details. (B) Same as (A) but for bimanual typing. Performance averaged over four testing blocks.

## Discussion

In today’s digital age, the ability to interact with devices such as computers and smartphones plays a crucial role in staying connected with friends and loved ones, as well as enabling access to the vast array of information available online. Unfortunately, individuals with tetraplegia, including those with locked-in syndrome (paralysis of almost all voluntary muscles), often find themselves cut off from these essential digital interactions. To bridge this gap, a variety of assistive communication technologies have been developed, from eye-tracking systems to advanced neural interfaces. These technologies have made significant strides, enabling control over 2D cursors (Bacher et al. 2015; Jarosiewicz et al. 2015; Gilja et al. 2015; Pandarinath et al. 2017), facilitating handwriting (Willett et al. 2021), and even supporting speech (Willett et al. 2023; Metzger et al. 2023; Card et al. 2023). Despite advancements in voice-to-text transcription, there is a lack of technology that enables users to type directly through finger movements. This technology would replicate the functionality of the widely used QWERTY keyboard, a tool that remains essential for computer use. We focused on developing a neural interface that enables typewriting by interpreting attempted finger movements. This approach aims to offer the same speed and flexibility that individuals experience when using a traditional computer keyboard.

In our study, we explored the adaptability of the typing interface through two distinct BCI paradigms. The first paradigm, point-and-click typing, utilizes real-time virtual finger movements to create a versatile interface. This method achieved a typing speed of 30-40 correct characters per minute for English sentences, matching the typing speed observed in traditional point-and-click methods using a 2D cursor (Pandarinath et al. 2017). The second paradigm, keystroke typing, involves a series of discrete finger movements without real-time feedback, resulting in notably higher speeds. When combined with effective error correction techniques to account for the uncertainty in character decoding, this approach allows for a balance between speed and accuracy, echoing the recent advancements in neural interface speeds for both handwriting and speech. Remarkably, typing English sentences via keystrokes more than doubled the speed achievable with point-and-click typing, matching the 90 characters per minute reported for handwriting decoding (Willett et al. 2021). While our evaluation primarily focused on cued character selection, with practice and familiarization with the keyboard layout, users could potentially approach these maximum performance levels.

Beyond these two paradigms, the flexibility of finger movements enables further adaptability and generalization of the interface. First, the interface can be used with one or two hands (with the current study being the first to enable closed-loop movements of bimanual fingers from a unilateral implant). Second, either one character can be selected at a time or multiple characters can be selected simultaneously with different fingers. Third, a shared decoder can be deployed across these typing strategies allowing users to quickly switch between them. Fourth, a hybrid of the point-and-click and keystroke paradigm is possible in the future – the likely sequence of characters can be identified by classifying finger trajectories produced using closed-loop control. By removing the slower aspects of closed-loop control such as the time spent clicking at the end of each trial, this approach could potentially improve the typing speed. Finally, the continuous velocity decoder for finger movements could be adapted to other tasks such as enabling a virtual joystick for video game control (Willsey et al. 2024). This level of adaptability empowers BCI users to customize the interface to their unique needs, potentially enhancing the likelihood of widespread adoption of this technology (Scherer et al. 2005; Blabe et al. 2015; Fried-Oken, Mooney, and Peters 2015; Pitt and Brumberg 2018).

The balance between flexibility and high performance is achieved through several key factors. First, finger movements offer multiple, largely independent degrees of freedom, unified under a shared structure of kinematics and neural representation. In able-bodied individuals, these distinctive properties of finger movements enable astonishing feats of precision and dexterity, such as intricate musical instrument performance, detailed artistry, and sophisticated sign language communication. For BCI users, finger movements provide an opportunity for a high degree of freedom control with reduced cognitive load.

Second, we use the statistics of the English language to simplify the finger movements required for typing. For the point-and-click mode, the high-frequency letters were placed closer to the starting position, thereby minimizing the average travel distance. For the keystroke sequences, the vocabulary of 31 characters was compressed into 12 distinct finger movements. While this compression is lossy, a language model was able to effectively reconstruct sentences with low errors.

Finally, we optimized the interface to improve performance with the limited SNR provided by existing neural interfaces. To this end, we limited each finger to one axis of motion (flexion-extension) and confined the movement to three finger groups per hand. We did not employ the standard QWERTY keyboard layout in this study, as it requires complex wrist and finger movements. Additionally, frequently used letters like E, T, and A are positioned on the side of the hand with lower SNR in our participant. However, we anticipate that future intracortical BCIs, perhaps incorporating increased channel counts, inputs from both brain hemispheres, and an advanced language model could potentially enable QWERTY typing.

Overall, this work introduces a high-performance typing intracortical BCI system that utilizes finger movements and offers customizable configurations spanning continuous and discrete movements, one or more fingers, and one or both hands. This work lays the foundation for future BCI designs that use a high degree of freedom movements to provide flexibility and prioritize end-user needs.

## Acknowledgment

The authors would like to thank T5 and his care partners. We would like to thank Beverly Davis, Kathy Tsou, and Sandrin Kosasih for administrative support.

Support provided by Office of Research and Development, Rehabilitation R&D Service, Department of Veterans Affairs (N2864C, A2295R); Wu Tsai Neurosciences Institute; Howard Hughes Medical Institute; Larry and Pamela Garlick; Samuel and Betsy Reeves; Sons Foundation Collaboration on the Global Brain 543045; NIDCD R01-DC014034, NIDCD U01-DC017844, NINDS UH2-NS095548, NINDS U01-NS098968.

* The contents do not represent the views of the Department of Veterans Affairs or the US Government. CAUTION: Investigational Device. Limited by Federal Law to Investigational Use

## Methods

### Clinical trial and participant

The participant, T5, was enrolled in the BrainGate2 Neural Interface System clinical trial (NCT00912041, registered June 3, 2009) with an investigational device exemption (IDE #G090003). This study was approved by the Institutional Review Board (IRB) of Stanford University (protocol #20804) and the Mass General Brigham IRB (protocol #2009P000505). The participant, T5, was a 69-year-old right-handed man with a C4 AIS C spinal cord injury. In August 2016, two 96-channel microelectrode arrays (Neuroport arrays with 1.5-mm electrode length; Blackrock Microsystems, Salt Lake City, UT) were placed in the anatomically identified hand ‘knob’ area of the left precentral gyrus. Detailed array locations are depicted on an MRI-reconstructed graphic in (Deo et al. 2023) in Extended Data Fig. 1a. Below the level of injury, T5 had very low amplitude movements of his limbs that consisted primarily of muscle twitching (Willett et al. 2020) for neurologic exam details.

### Neural recordings

The BCI rig was set up in two distinct configurations as our lab transitioned from an older analog setup to a newer digital setup. In the first setup used until 7/10/2023, the BCI rig was set up with 2 patient cables connected to transcutaneous pedestals, which were routed to the Neural Signal Front End Amplifier (Blackrock Neurotech, Salt Lake City, Utah) where the raw voltage was bandpass filtered (0.3 Hz first-order high-pass and 7.5 kHz third-order low-pass), sampled at 30 kHz with 250 nV resolution, converted to an optical signal, and then sent to the Neural Signal Processor (Microsystems 2023). From 7/26/2023 and later, 2 Neuroplex E headstages were connected to 2 transcutaneous pedestals, and the signal was analog filtered and sampled at the headstage and then sent to the digital hub via a micro-HDMI cable. At the digital hub, the signal was converted to an optical signal transmitted via optical cable to the Neural Signal Processor. The Neural Signal Processor sent the digital signal to a SuperLogics machine running Simulink Real-Time (v2019, Mathworks, Natick, MA). Common average referencing (CAR) was used to reduce the electrical noise on the input channels for ‘point-and-click’ sessions, which were performed before June 2023. Due to additional noise in recordings after June 2023, linear regression referencing (Young et al. 2018) was used for ‘keystroke’ typing sessions.

For threshold crossing detection, we used a −4.5 x RMS threshold applied to each electrode, where RMS is the electrode-specific root mean square (standard deviation) of the voltage-time series recorded on that electrode. Threshold crossing counts were computed in 20 ms bins for analysis and decoding. To compute spike-band power (SBP), the signals then passed through a 250-Hz digital high-pass filter, squared and summed in 20-ms windows.

Every 20 ms, UDP packets of neural features were communicated to a Linux computer running Ubuntu with Python (v3.7.11), Tensorflow 2.7 (https://www.tensorflow.org/), and Redis (v7.02), which implemented the decoding algorithm. The entire system was interfaced with an additional Windows computer running Matlab (v2019, Mathworks, Natick, MA) that was used to stop and start experimental runs during sessions.

### Keyboard design

A virtual keyboard with a finger and hand visualization was developed in Unity Software (Unity Technologies, San Francisco) using a pre-made hand model (Barnett, n.d.). The fingers and hand were animated by specifying the trajectory of joint positions and rotations directly from an external program using Redis. Finger motion involved only the flexion-extension movements of individual fingers and joint positions and angles were interpolated between a complete flexion and extension position.

Keys were evenly spaced between complete flexion (=0) and complete extension (=1, spacing = 0.15 units), with the neutral rest corresponding to 0.5. Keys on nearby fingers were staggered with respect to each other so that unique characters could still be selected when fingers were constrained to move together as a single finger group (e.g. index/middle and ring/little).

Variants of the keyboard were developed for either using the right hand only or using both hands (bimanual keyboard). Characters were assigned to keys based on the unigram frequencies of characters in the English language. Thirty-one characters (26 letters + four symbols) were considered, matching the character set used in (Willett et al. 2021). Unigram frequencies were estimated using 57340 sentences in the Brown Corpus (Francis and Kucera 1979). High-frequency symbols were placed closer to rest and evenly distributed across finger groups. Characters were assigned to keys greedily in order of decreasing frequency, with a preference for flexion over extension, right hand over left hand (for the bimanual keyboard), and thumb over index-middle and index-middle over ring-little finger groups.

### Point-and-click character selection

Point-and-click typing consists of moving a virtual finger over a key followed by selecting it with a ‘click’, triggered by a rapid attempted movement of another (non-finger) effector. Typing was evaluated with cued character selection since free typing would require memorizing the location of all symbols, requiring substantial practice. Character sequences forming sentences from (Willett et al. 2021) were cued.

The target characters were cued (in red) at the beginning of each trial and started with fingers at rest (midway between complete flexion and extension). As the participant attempted to move the virtual fingers, the selected character was highlighted in blue. The participant initiates a click to finalize the selection. Trials finish as soon as a click is administered, and the character indicated by the fingers is selected. Finger positions were then reset to the rest position before the beginning of the next trial.

Clicks were decoded using a logistic regression-based classifier. The click-classifier was learned using an open-loop finger movement task, where each trial ended in an attempted click. A click was administered if the click probability was above a threshold of 0.5 continuously for a hold period (30-80ms). A refractory period (450ms) was administered after the click, where clicks could not be re-administered.

A neural-network-based decoder estimated finger velocities from neural activity recordings (see below for details). The finger velocities were used to update finger position only if the click probability was below a threshold of 0.5.

Three configurations of the point-and-click character selection were tested: (1) selecting one character in a sentence per trial; using the right-hand keyboard with three finger groups; clicking using the ‘jerking’ movement of the left-hand elbow, (2) selecting one character per trial; using the bimanual keyboard with three finger groups each; clicking using the movement of both feet and (3) selecting up to two characters per trial if successive characters in a sentence are on different finger groups; using the right-hand keyboard with three finger groups; clicking using the movement of the left-hand elbow (same as 1).

For selecting characters after a click, the fingers sufficiently far from their rest position are first identified and the closest one (or two, depending on the task) character(s) to the corresponding non-rest finger are selected.

Note that, when multiple characters are cued per trial, a language model is necessary to identify the order of selected characters. The details of language model usage are described below.

### Continuous velocity decoding for point-and-click typing

We developed a recipe for estimating the parameters of a non-linear decoder for continuous control of multiple degrees of freedom movements. We describe four steps for building the decoder pipeline, each designed to solve a unique challenge associated with a high degree of freedom motor decoding. These steps consist of (1) collecting training data (addresses distribution shift in neural activity between open-loop and closed-loop movements), (2) deciding decoder architecture (addresses non-linear representation of movements), (3) training the decoder (reduces data required for training neural network decoders) and (4) postprocessing of predicted velocities (incorporates task-related priors).

#### 1. Training data gathered during closed-loop movements

The first problem is the difference in the neural activity between open-loop movements (i.e., without instantaneous visual feedback), which is typically used to build the initial decoder, and closed-loop movements (i.e., with instantaneous visual feedback), which is needed for typing (Koyama et al. 2010; Jarosiewicz et al. 2013). As shown in Fig. S2a, mean activity across channels was different between nearby blocks of open-loop and closed-loop movements. Since the eventual decoder usage will be during closed-loop movements, we aim to maximize the amount of closed-loop data used for training. One way is to deploy the decoder in closed-loop and continuously update the decoder as more training data is collected (Fig. 2C, (Brandman et al. 2018)).

For the finger movement decoding considered in this paper, training data was collected during a point-and-hold task that cues a single finger group movement in each trial. Target keys were randomly cued with uniform probability across finger directions. The participant attempts to move the finger corresponding to the target key with a logistic regression-based classifier. The classifier either predicts no movement or the movement of one finger group. For the predicted finger group, the position is updated with a constant velocity. When the total time spent over a key exceeds 2 seconds during a trial, the key is selected and the next trial begins.

At each time step, the training data is augmented by logging the neural activity and corresponding attempted movements (assuming that only the target finger is moving towards the target key and the other fingers are at rest), and the logistic-regression-based classifier is learned from scratch (implemented in sklearn; Pedregosa et al. 2011) continuously.

Note that the decoder is not available until all movements have been attempted at least once, necessitating that a small (∼10) number of initial trials are open-loop. At the end of a five-minute long block with nearly 100 trials, the training set consists of mostly closed-loop data.

#### 2. Movement decoding using a neural network model

The second problem is the non-linear representation of finger movements - the magnitude of neural activity is non-linearly related to the finger velocities (Willsey et al. 2022; Temmar et al. 2024), and the activities of individual fingers sum pseudo-linearly during simultaneous finger movements (Shah et al. 2023). This calls for using a non-linear model to decode attempted finger velocities from neural activity. We use a fully connected, feed-forward neural network that relates the instantaneous neural activity across multiple-time steps (5 steps of 20ms bins resulting in 100 total ms of neural activity) to the attempted velocities (Fig. 2A). The input is a concatenation of neural activity (threshold crossings, with SBP included for later sessions, see Table S1, S2, and S3). The decoder architecture has a concatenation of four modules, with each module consisting of a fully connected neural network, or dense layer (256 units), dropout (with rate 0.1), batch normalization and rectifying nonlinearity (f(x) = max(x, 0)). The architecture is similar to those previously reported (Willsey et al. 2022, 2024) although without the initial convolutional layer, which reduces the number of learnable parameters; using five 20ms time-bins of threshold crossing and SBP instead of three 50ms time-bins of SBP. The output of the decoder was velocities for each finger group (three outputs for the right-hand keyboard and six outputs for the bimanual keyboard). The attempted movements were used as output labels for training and were labeled based on the relative location of each finger compared to the target, either as +1 (extension), 0 (rest), or −1 (flexion). The decoder was trained with mean-squared error loss.

The decoder was implemented in Tensorflow 2.7 and trained with Adam optimizer (step size 0.0001, batch size 200). Data was partitioned into training (80%) and evaluation data (20%), and training was stopped when the loss on evaluation data (calculated every 20 training steps) did not decrease in the last 200 evaluation steps.

#### 3. Pre-training and fine-tuning to reduce in-session calibration time

The third problem is that neural network decoders with a large number of learned parameters typically require a large amount of training data, which may be more than the amount that can be recorded in a single 5-minute block. To reduce the calibration times, the decoder was pretrained on the data collected from previous sessions and fine-tuned on the data collected on a target session day. Both pretraining and fine-tuning used the closed-loop data collected with a logistic-regression classifier-based decoder updating in real-time, as described in the ‘Training data’ section above. The pre-training and fine-tuning approach improved performance compared to training the model from scratch (i.e., random initialization of decoder parameters, Fig. 2D).

#### 4. Post-processing to reduce finger drift

The fourth problem is that the errors in finger velocity predictions can accumulate over timesteps, resulting in drifting finger positions. Specifically, when the target velocities are zero (i.e., fingers are not supposed to move), the trained decoder typically predicts low amplitude values dispersed around zero, and this velocity is integrated over multiple time steps to result in a drift for the finger. The drift needs a participant-initiated corrective movement back to rest, making the fingers less controllable. This issue was minimized with soft-thresholding, which makes all predictions less than a certain threshold around rest exactly zero and thus removing drift (Fig. 2E). In A-B-A-B testing, incorporating soft-thresholding was shown to improve performance (Supp. Fig. 3). In addition to soft-thresholding, other post-processing steps including applying a hand-tuned scalar gain to the decoded velocities (identical across fingers), smoothening the velocities across successive time steps, and applying a power function (*f(x) = x^p^*, p>1) to account for a non-linear relationship between attempted movement strength and neural activity magnitude as shown in (Willsey et al. 2022)). The hyperparameters were selected by manual quasi-optimization and are detailed in Tables S1, S2, and S3.

### Keystroke character selection

Keystroke character selection was evaluated using the bimanual keyboard with three finger groups on each hand. A letter was cued at regular intervals (1sec, 0.8, or 0.67 sec) and the participant was instructed to attempt movement of the corresponding finger in the corresponding direction. No closed-loop feedback was given during the attempted movements.

Selected character sequences were estimated by first decoding finger-group movements from neural recordings and followed by error correction using a language model. A logistic regression classifier decoded the summed neural activity (both threshold crossings and Spike Band Power after Z-scoring within each block) within a trial into one of 12 different finger movement directions (finger movement classes correspond to all combinations of flexion/extension, thumb/index-middle/ring-little finger groups, and left/right hands). The classifier was tested on all trials of a sentence after training it on all other sentences. Hence, the classifier gives an array of finger movement probabilities for each trial. For faster speeds (0.67sec / trial), the analyzed window of neural activity was delayed by 0.25 seconds from cue onset.

This array was converted into an array of character probabilities, by applying a uniform distribution to all characters along a finger movement direction in the bimanual keyboard. For example, as ‘T’, ‘M’, and ‘X’ all correspond to left-hand thumb extension, *p(character ‘T’) = (1/3) x p (left thumb extension)* for the trial.

### Language model

The details of the language model (LM) are described in (Fan et al. 2023). Briefly, a 3-gram LM, along with a beam search algorithm, processed a character probability output and translated it into words. The 3-gram LM was trained on the OpenWebText2 (Gao et al. 2020) corpus using Kaldi (Povey, Ghoshal, and Boulianne 2011). It uses a 130,000-word vocabulary taken from the CMU Pronouncing Dictionary (“The CMU Pronouncing Dictionary,” n.d.). Below, we describe how the character probability output was constructed for (i) point-and-click typing with two character selections per trial and (ii) keystroke typing.

#### Bimanual Keystroke character selection

For each keystroke trial, a sequence of two probability vectors was constructed to use as input to the language model, each of length N+1, where N is the size of the character set (N=31 in this work). The extra symbol encodes a ‘blank’ symbol used for Connectionist Temporal Classification (CTC) loss (Graves et al. 2006) and indicates the beginning of a new character in the sequence. The first vector corresponds to the probability of different characters described above (with 0 probability for the blank symbol). The second vector corresponds to zero probability for all symbols other than the blank symbol. Hence, a character sequence with L characters and character set size N (=31 in this work) is converted to an array of probabilities of size 2L x (N + 1). The LM evaluates this array to give the most likely English sentence.

#### Point-and-click character selection, up to two characters per trial, right-hand keyboard

In this case, the primary role of the LM is to identify the order of characters that are selected simultaneously in each trial. Additionally, it could also reduce the error from incorrect character selections - namely, in cases where the wrong number of characters is selected (one character selected instead of two target characters, or two characters instead of one); or the wrong set of characters is selected even if the number of selected characters is correct. To account for all these possibilities, the selected characters were converted to an array of probability vectors with each trial corresponding to six probability vectors. The first and third vectors encode the selected characters, with the log probability of selected characters being 0, other characters being −5, and blank symbols being −5. The second and fourth vectors allow for an additional character with the log probability of all characters being −3 and the blank symbol being 0. The third and sixth vectors allow for the blank symbol with the log probability of all characters being −5 and the blank symbol being 0.

### Performance metrics

Typing performance was characterized by measuring the accuracy and speed of character selection. The success rate measures the fraction of correct character selections across all the trials in a block. Correct Characters Per Minute (CCPM) for a block is computed as # correct character selections / total time. Character error rate is calculated by summing the edit distances between the target sentence and the decoded sentence for all target sentences within a block, and then dividing by the total length of all target sentences. Edit distance is calculated as the minimum number of additions, deletions, or substitutions required to transform one string into the other.

Decoding accuracy is measured using the alignment of the predicted velocity to the target velocity at each time step. Directional signal-to-noise Ratio (dSNR) is the ratio of the component of velocity along the target velocity direction (i.e. signal) and the component orthogonal to the target velocity direction (i.e. noise). Mathematically, for the target velocity unit vector *t̂_i_*, predicted velocity *p_i_* at the time step *i*, dSNR is given by (see (Willsey et al. 2024) for further details):

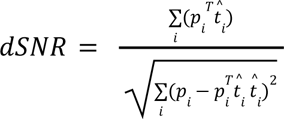

### Multi-task training

For the multi-task training analysis depicted in Fig. 5, the neural network decoder described above was used, with a modification to allow linearly reading out the continuous output (finger velocity) and discrete output (finger class) from the last layer. Both discrete and continuous outputs were used for each task (either point-and-hold or keystroke typing). The continuous output was trained with mean squared error to velocity output (+1 for extension, −1 for flexion, and 0 for no movement). The discrete output was trained with the cross-entropy loss to the discrete movement classes (13 classes, with 12 classes corresponding to finger movement combinations of flexion/extension movements on three-finger groups on either hand and one class for no movement). When applying the decoder on the right-hand tasks, the velocity output for the left hand was trained to be zero (as the participant was not attempting to move the left hand).

Only the continuous output was used for evaluation, as it is the only task-relevant output for the point-and-hold task. For the right-hand keyboard, only the velocity output for the right hand was used.

For pretraining, 11 training blocks (with one block per session) for right-hand point-and-click (closed-loop data), 7 blocks for bimanual point-and-click (closed-loop data), and 17 blocks for keystroke typing (open-loop data) were used. Fine-tuning was performed on increasing amounts of training data from a block and evaluated on held-out data from the same block. For fine-tuning and evaluation, ten blocks for right-hand keyboard point-and-hold and 4 blocks for bimanual keyboard point-and-hold were used. The final performance was averaged across all blocks.

## Supplementary information

**Fig. S1.**
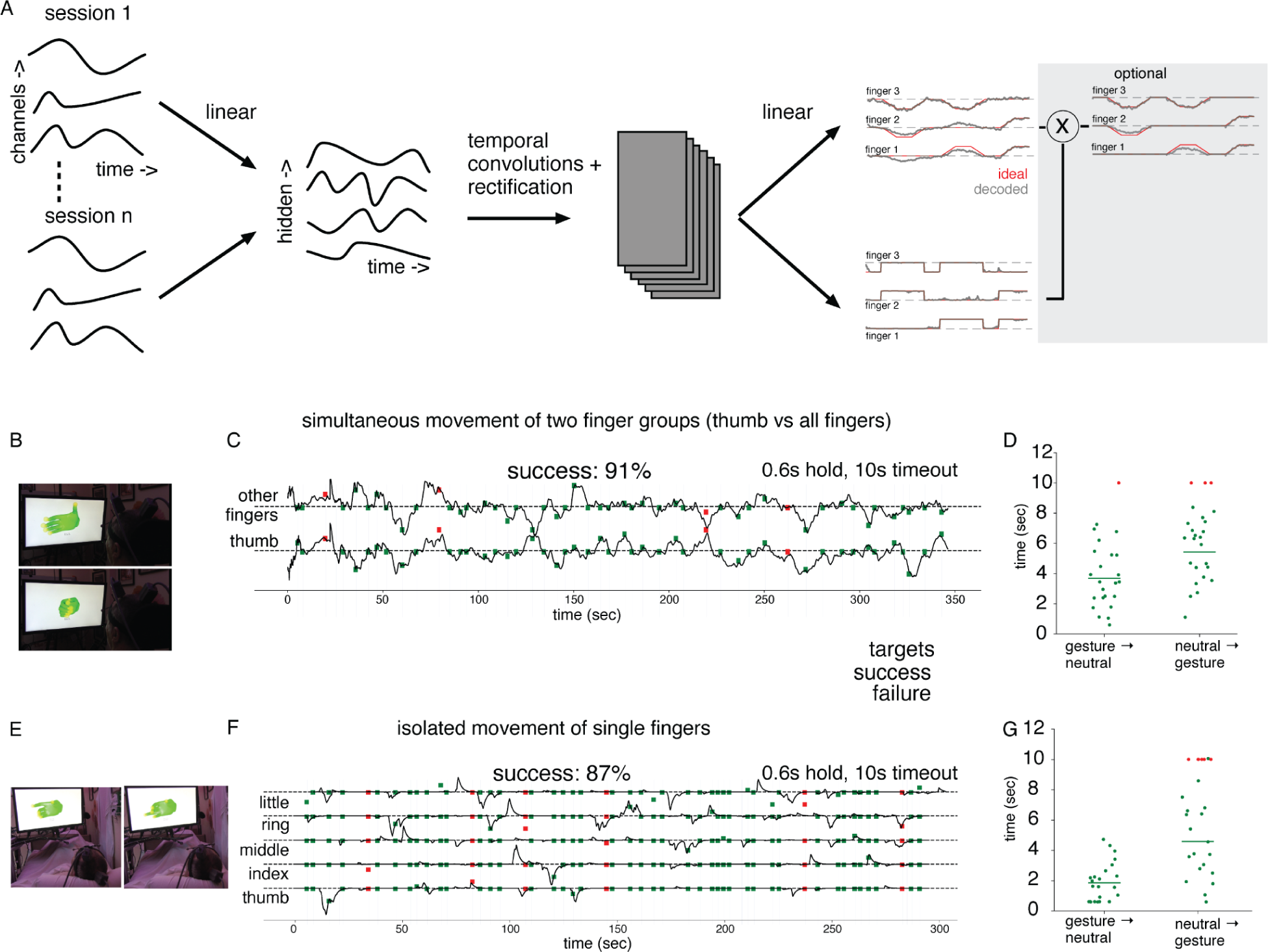
Closed-loop position control of finger movements. (A) Decoder architecture: Binned multi-unit neural activity is passed through a day-specific linear filter, followed by a temporal convolutional layer and rectification. The two separate linear readouts predict the positions of individual fingers (continuous output) and movement probability (either flexion or extension) of individual fingers (discrete output) respectively. For the isolated single-finger movement task, the continuous finger position output is used only if the discrete finger movement probability is above a threshold. (B, C, D) Simultaneous movement of two finger groups (thumb, and all other fingers linked together). (B) Examples of achieved gestures. (C) Decoded finger positions for a single block. Squares indicate target positions for two finger groups. Each trial consisted of a preparatory period (where the target was indicated but movement was not required) and a move period (where T5 attempted to move toward the target). Trials alternated between going from neutral/rest position to a random target position and going back to neutral/rest. Target positions were sampled independently between 0-1, with a grid size of 0.1; the rest corresponds to 0.5 Trials were successful (green squares) if both finger groups were within 10% of the target position for both fingers for 0.6s continuously, and are declared a failure (red squares) if the target was not attained after 10s. (D) Target acquisition times, separated for the two types of trials: going out to a target gesture (neutral → gesture) and going back to the neutral gesture (gesture → neutral). Each dot is a single trial, successful trials are shown in green, and failed trials are shown in red. (E, F, G) Same as (B-D) for isolated movements of single fingers, where one (and only one) finger is required to move in each trial.

**Fig. S2.**
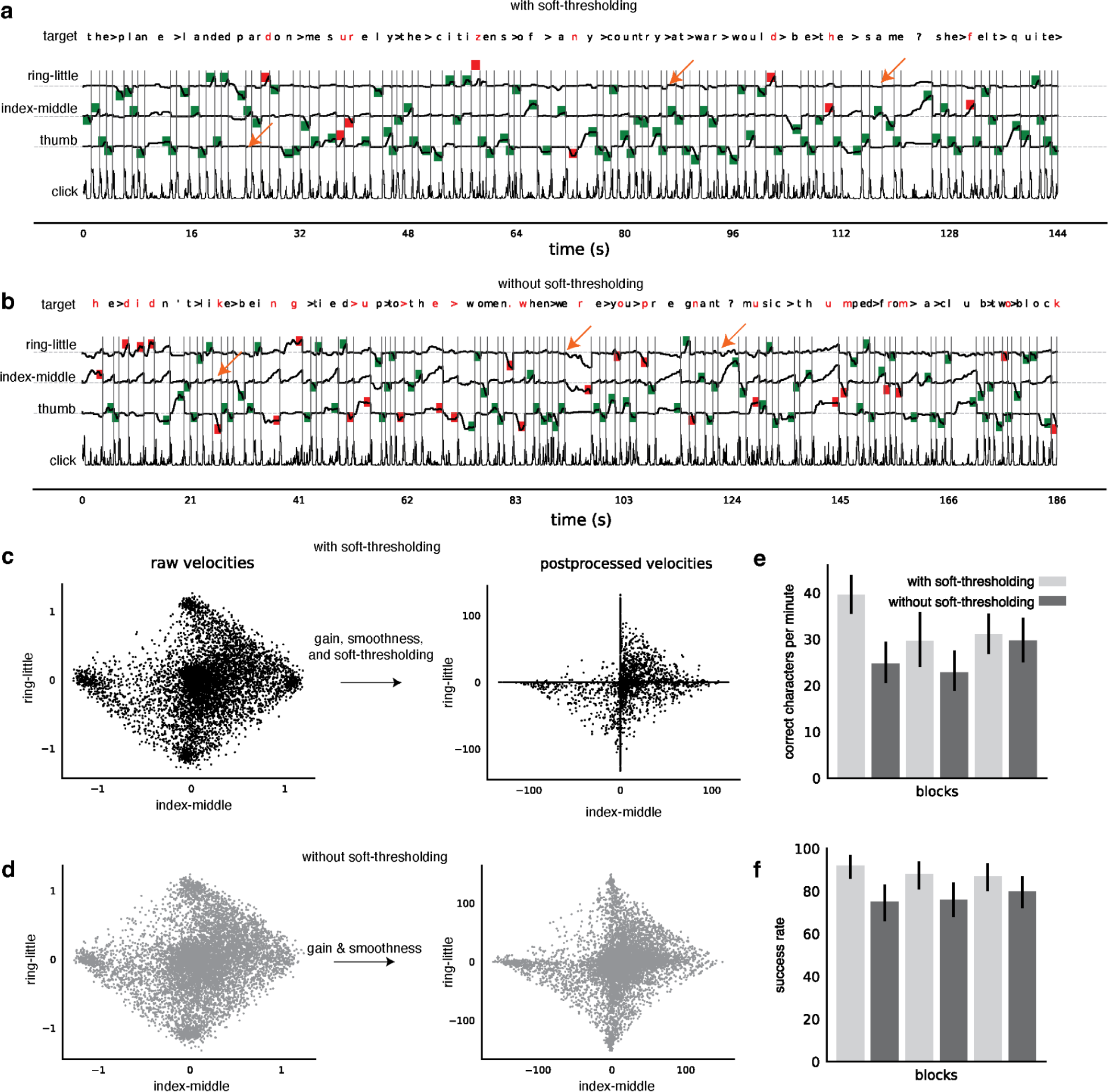
Soft-thresholding of predicted velocities for multiple degrees of freedom decoding. (A, B) Soft-thresholding function, given by f(x) = max(x-d, 0) - max(-x - d, 0) zeros out any x between -d and d, and reduces the magnitude by d for any |x| > d. Finger trajectories in a block with soft-thresholding used (A) or not used (B). Note the jittering positions of non-target fingers (indicated by arrows) when soft-thresholding was not used, which is absent when sof-threshold is used. (C) reproduced from Fig. 3D. (D) similar to (C), without soft-thresholding. (E, F) A-B-A-B testing of closed-loop finger control with and without soft-thresholding. Performance was measured by the correct character selections per minute (E) and accuracy (F). 95% Confidence intervals computed by bootstrapping (10000 resamplings). For both the measures, the difference between all trials with soft-thresholding and without soft-thresholding was significant (p<0.001, bootstrapping with 10000 resamplings).

**Fig. S3.**
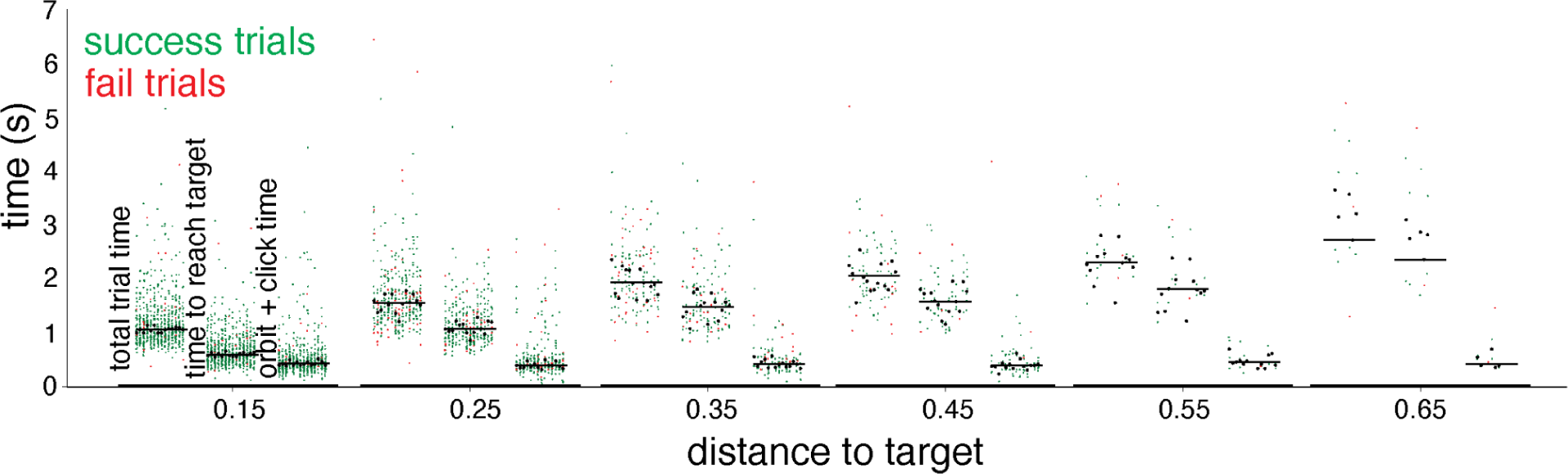
Trial times for point-and-click on the right-hand keyboard with one character per trial. Trial completion times for the right-hand keyboard (one character per trial), separated by the distance of the target key from the rest (x-axis). Total trial times, time to first reach the target, and the orbit + click time (measured as the difference between total trial time and first reaching the target) are separated. Successful (failed) trials are indicated by green dots (red). Trials can fail due to a time-out of 10 seconds or incorrect character selection. For each distance, trials are grouped by the recording block, with the within-block mean indicated by black dots and the mean across all blocks indicated by the black line. Note that the number of target keys decreases with increasing distance (as intended by the keyboard design); total trial time and time to target increases with the distance of the target key and orbit + click time does not change with the distance of the target key.

**Fig. S4.**
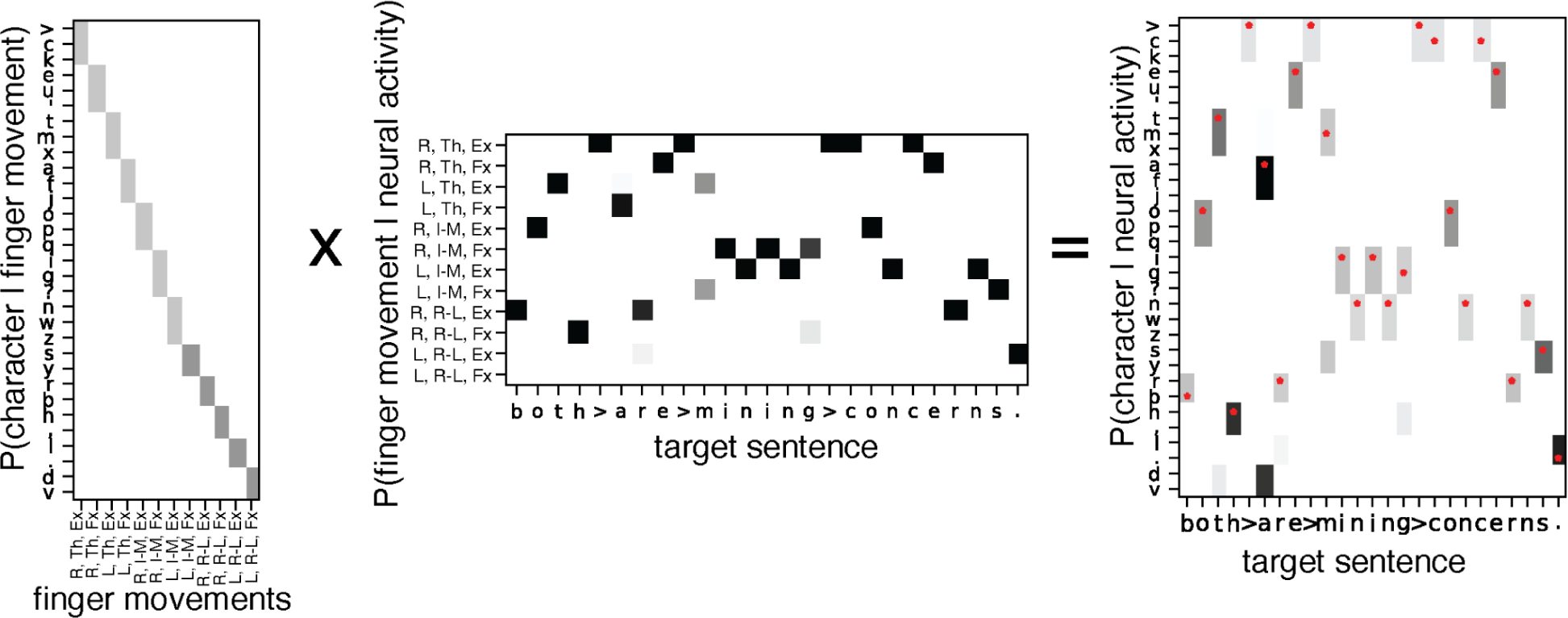
Details of neural decoding for keystroke typing using the bimanual keyboard. A decoder uses neural activity recorded concurrently with the finger movements and predicts a probability distribution over finger movements, which is then converted into a distribution over characters assuming the equal probability of different characters along a finger movement direction (ex. left thumb extension corresponds to a uniform distribution over symbols T, M and X). The probability distribution associated with each character in a sentence is then passed through a language model. Finger movements are indicated by a tuple of laterality (L/R), finger group (thumb: T, Index-Middle: I-M, Ring-Little: R-L), and movement (Extension: Ex, Flexion: Fx).

**Video S1. Real-time position control of two finger groups.** Recording of the closed-loop BCI control of simultaneous movements of two finger groups. Multiple trials are shown in succession, with each trial consisting of a preparatory period (red hands) and a move period (green hands). Target positions at the end of the trial are shown with large markers and small markers indicate the current finger positions. T5 was instructed to observe and prepare for the target (but not move) during the preparatory period. Trials are successful if the markers at the end of each finger are close (turn yellow) to the target positions for the corresponding finger. Target positions consist of different degrees of flexion and extension of the two finger groups, sampled independently.

**Video S2 Real-time position control of isolated movements of individual fingers.** Similar to Video S1, but for isolated movements of individual fingers.

**Video S3. Right Hand ‘Point-and-Click’ Typing Using Three Finger Groups.** Recording of 100 trials of closed-loop BCI control for typing. Each trial begins with a cued character, marked in red, that forms part of an English sentence. As T5 directs the relevant finger toward the cued key, the key being selected is indicated in blue. Selection is confirmed through ‘clicking’, which corresponds to a swift left elbow movement. The completion of a trial is marked by either a successful selection of the cued key or a failure, characterized by the selection of an uncued key or no selection within a 10-second window; each outcome is audibly distinguished. The video includes an on-screen display of the target sentence (top row), the sentence decoded from the participant’s selections (second row), and corresponding performance metrics (third row) for accuracy (percent of successful character selections) and speed (number of correct characters per minute).

**Video S4. Bimanual ‘Point-and-Click Typing’ Using Three Finger Groups.** Similar to Video S3, but with a bimanual keyboard and clicking corresponding to swift gas-pedal movement of ankles on both feet.

**Video S5. Right-hand ‘Point-and-click’ typing with three finger groups and a selection of up to two characters per trial.** If two successive characters in a sentence lie on different finger groups, they are cued in the same trial. T5 selects both the cued characters before initiating a click. Trials are successful only if all the cued characters are selected. The video includes an on-screen display of the target sentence (top row), the sentence decoded from the participant’s selections assuming a perfect language model (second row), and corresponding performance metrics for number of correct character selections (third row). Note that the results in the main text do not assume a perfect language model. Other details are the same as in Video S3.

**Table S1.** Parameters for “Point-and-Click” typing using the right hand, three finger groups, one target/trial, click with the left hand. Tx indicated threshold crossings and SBP indicates Spike Band Power.

| Dataset | Neural features used | Velocity Gain | Velocity Smoothness | Velocity Soft thresholding | Velocity power | Click threshold | Click hold duration | Click blankout | Accuracy (%) | CCP M |
| --- | --- | --- | --- | --- | --- | --- | --- | --- | --- | --- |
| t5.2023.01.03, block 5 | TX | 80 | 5 | 0 | 1 | 0.5 | 150ms | 450ms | 92 | 30.9 |
| t5.2023.01.03, block 6 | TX | 80 | 5 | 0.1 | 1 | 0.5 | 150ms | 450ms | 96 | 33.2 |
| t5.2023.01.03, block 7 | TX | 80 | 5 | 0.1 | 1 | 0.5 | 150ms | 450ms | 95 | 30.1 |
| T5.2023.01.17, block12 | Tx | 80 | 5 | 0.1 | 1 | 0.5 | 80ms | 450ms | 90 | 29.7 |
| T5.2023.01.17, block 13 | Tx | 80 | 5 | 0.1 | 1 | 0.5 | 80ms | 450ms | 87 | 31.1 |
| T5.2023.01.17, block 14 | Tx | 80 | 5 | 0.1 | 1 | 0.5 | 80ms | 450ms | 85 | 34.0 |
| T5.2023.01.17, block 15 | Tx | 80 | 5 | 0.1 | 1 | 0.5 | 80ms | 450ms | 87 | 30.1 |
| T5.2023.01.19, block 4 | Tx | 80 | 5 | 0.1 | 1 | 0.5 | 80ms | 450ms | 87 | 30.0 |
| T5.2023.01.19, block 5 | Tx | 100 | 5 | 0.12 | 1 | 0.5 | 80ms | 450ms | 87 | 32.4 |
| T5.2023.01.19, block 6 | Tx | 100 | 5 | 0.12 | 1 | 0.5 | 80ms | 450ms | 87 | 30.8 |
| T5.2023.01.19, block 19 | Tx | 100 | 5 | 0.12 | 1 | 0.5 | 80ms | 450ms | 88 | 33.7 |
| t5.2023.05.03, block 20 | Tx + SBP | 125 | 5 | 0.16 | 1.3 | 0.5 | 40ms | 450ms | 92 | 38.4 |
| t5.2023.05.03, block 22 | Tx + SBP | 125 | 5 | 0.16 | 1.3 | 0.5 | 40ms | 450ms | 88 | 28.8 |
| t5.2023.05.03, block 24 | Tx + SBP | 125 | 5 | 0.16 | 1.3 | 0.5 | 40ms | 450ms | 87 | 30.3 |

**Table S2.**
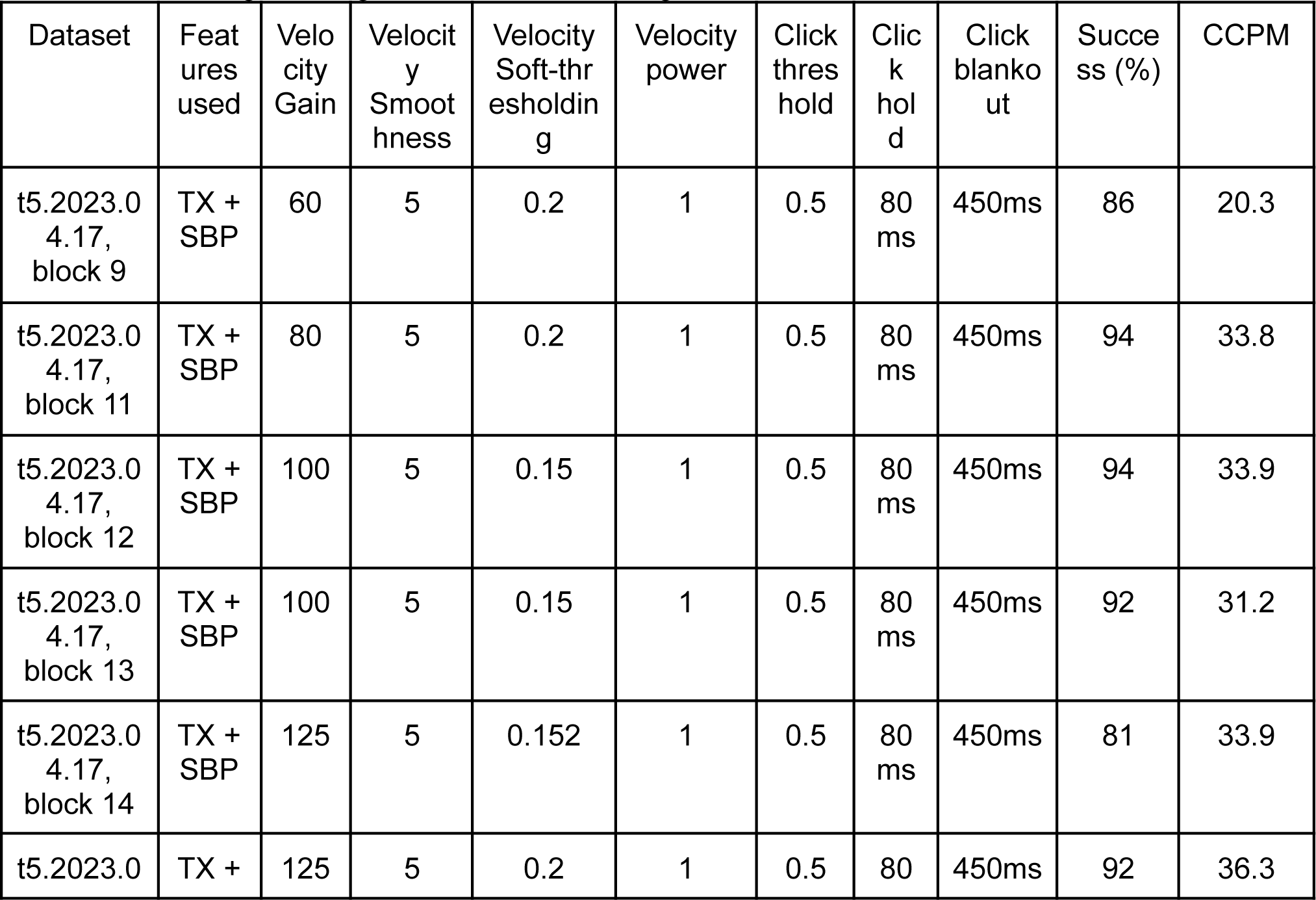
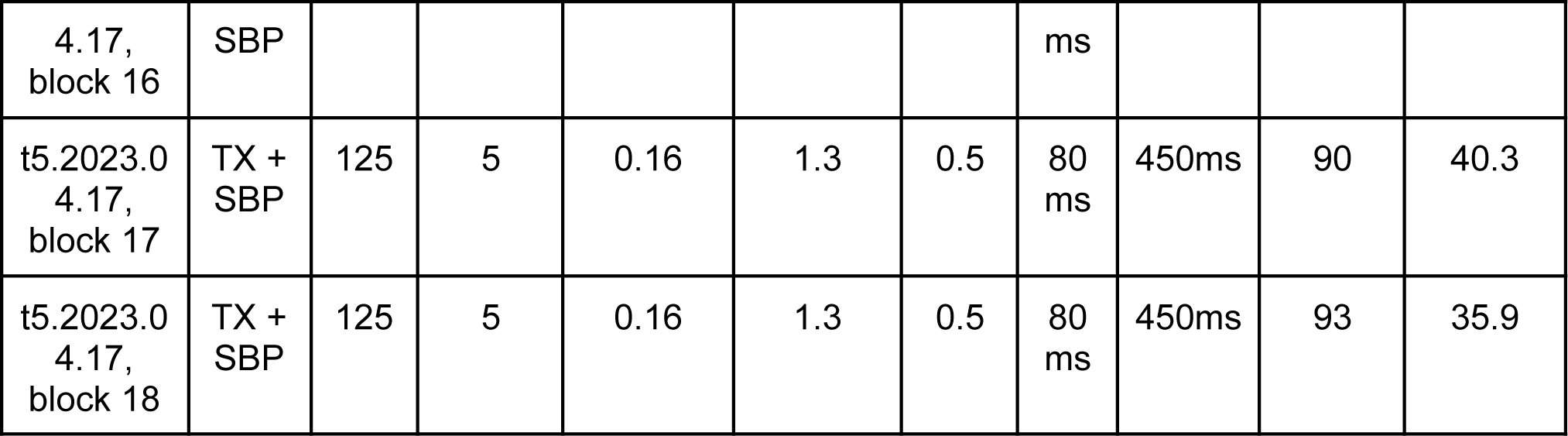
Parameters for “Point-and-Click” typing using the right hand, three finger groups, two chars / trial, moving one finger at a time, click using the left hand.

**Table S3.** Parameters for “Point-and-Click” typing using both hands (bimanual), three finger groups on each hand, one target/trial, and click with both feet.

| Dataset | Features used | Velocity Gain | Velocity Smoothness | Velocity Soft-thresholding | Velocity power | Click threshold | Click hold (ms) | Click blankout (ms) | Success (%) | CCPM |
| --- | --- | --- | --- | --- | --- | --- | --- | --- | --- | --- |
| t5.2023.05<br>.22 block 17 | Tx+SBP | 125 | 5 | 0.16 | 1.3 | 0.5 | 30ms | 450ms | 87 | 24.2 |
| t5.2023.05<br>.22 block 18 | Tx+SBP | 125 | 5 | 0.16 | 1.3 | 0.5 | 30ms | 450ms | 87 | 30.5 |
| t5.2023.05<br>.22 block 19 | Tx+SBP | 125 | 5 | 0.16 | 1.3 | 0.5 | 30ms | 450ms | 90 | 25.4 |
| t5.2023.05<br>.22 block 21 | Tx+SBP | 125 | 5 | 0.16 | 1.3 | 0.5 | 30ms | 450ms | 83 | 25.7 |

## References

1. Anumanchipalli, Gopala K., Josh Chartier, and Edward F. Chang. 2019. “Speech Synthesis from Neural Decoding of Spoken Sentences.” Nature 568 (7753): 493–98.

2. Bacher, Daniel, Beata Jarosiewicz, Nicolas Y. Masse, Sergey D. Stavisky, John D. Simeral, Katherine Newell, Erin M. Oakley, Sydney S. Cash, Gerhard Friehs, and Leigh R. Hochberg. 2015. “Neural Point-and-Click Communication by a Person With Incomplete Locked-In Syndrome.” Neurorehabilitation and Neural Repair 29 (5): 462–71.

3. Barnett, Justin P. n.d. “Oculus VR Hands Models.” https://www.patreon.com/posts/free-oculus-vr-46544401.

4. Blabe, Christine H., Vikash Gilja, Cindy A. Chestek, Krishna V. Shenoy, Kim D. Anderson, and Jaimie M. Henderson. 2015. “Assessment of Brain–machine Interfaces from the Perspective of People with Paralysis.” Journal of Neural Engineering 12 (4): 043002.

5. Bommasani, Rishi, Drew A. Hudson, Ehsan Adeli, Russ Altman, Simran Arora, Sydney von Arx, Michael S. Bernstein, et al. 2021. “On the Opportunities and Risks of Foundation Models.” arXiv [cs.LG]. arXiv. http://arxiv.org/abs/2108.07258.

6. Brandman, David M., Tommy Hosman, Jad Saab, Michael C. Burkhart, Benjamin E. Shanahan, John G. Ciancibello, Anish A. Sarma, et al. 2018. “Rapid Calibration of an Intracortical Brain-Computer Interface for People with Tetraplegia.” Journal of Neural Engineering 15 (2): 026007.

7. Card, Nicholas S., Maitreyee Wairagkar, Carrina Iacobacci, Xianda Hou, Tyler Singer-Clark, Francis R. Willett, Erin M. Kunz, et al. 2023. “An Accurate and Rapidly Calibrating Speech Neuroprosthesis.” bioRxiv. 10.1101/2023.12.26.23300110.

8. Costello, Joseph T., Hisham Temmar, Luis H. Cubillos, Matthew J. Mender, Dylan M. Wallace, Matthew S. Willsey, Parag G. Patil, and Cynthia A. Chestek. 2023. “Balancing Memorization and Generalization in RNNs for High Performance Brain-Machine Interfaces.” *bioRxiv : The Preprint Server for Biology*, May. 10.1101/2023.05.28.542435.

9. Deo, Darrel R., Francis R. Willett, Donald T. Avansino, Leigh R. Hochberg, Jaimie M. Henderson, and Krishna V. Shenoy. 2023. “Translating Deep Learning to Neuroprosthetic Control.” *bioRxiv : The Preprint Server for Biology*, April. 10.1101/2023.04.21.537581.

10. Fan, Chaofei, Nick Hahn, Foram Kamdar, Donald Avansino, Guy H. Wilson, Leigh Hochberg, Krishna V. Shenoy, Jaimie M. Henderson, and Francis R. Willett. 2023. “Plug-and-Play Stability for Intracortical Brain-Computer Interfaces: A One-Year Demonstration of Seamless Brain-to-Text Communication.” ArXiv, November. https://www.ncbi.nlm.nih.gov/pubmed/37986728.

11. Francis, W. Nelson, and Henry Kucera. 1979. “Brown Corpus Manual.” Letters to the Editor 5 (2): 7.

12. Fried-Oken, Melanie, Aimee Mooney, and Betts Peters. 2015. “Supporting Communication for Patients with Neurodegenerative Disease.” NeuroRehabilitation 37 (1): 69–87.

13. Gao, Leo, Stella Biderman, Sid Black, Laurence Golding, Travis Hoppe, Charles Foster, Jason Phang, et al. 2020. “The Pile: An 800GB Dataset of Diverse Text for Language Modeling.” arXiv [cs.CL]. arXiv. http://arxiv.org/abs/2101.00027.

14. Gilja, Vikash, Paul Nuyujukian, Cindy A. Chestek, John P. Cunningham, Byron M. Yu, Joline M. Fan, Mark M. Churchland, et al. 2012. “A High-Performance Neural Prosthesis Enabled by Control Algorithm Design.” Nature Neuroscience 15 (12): 1752–57.

15. Gilja, Vikash, Chethan Pandarinath, Christine H. Blabe, Paul Nuyujukian, John D. Simeral, Anish A. Sarma, Brittany L. Sorice, et al. 2015. “Clinical Translation of a High-Performance Neural Prosthesis.” Nature Medicine 21 (10): 1142–45.

16. Graves, Alex, Santiago Fernández, Faustino Gomez, and Jürgen Schmidhuber. 2006. “Connectionist Temporal Classification: Labelling Unsegmented Sequence Data with Recurrent Neural Networks.” In Proceedings of the 23rd International Conference on Machine Learning, 369–76. ICML ‘06. New York, NY, USA: Association for Computing Machinery.

17. Guan, Charles, Tyson Aflalo, Kelly Kadlec, Jorge Gámez de Leon, Emily R. Rosario, Ausaf Bari, Nader Pouratian, and Richard A. Andersen. 2023. “Decoding and Geometry of Ten Finger Movements in Human Posterior Parietal Cortex and Motor Cortex.” Journal of Neural Engineering 20 (3). 10.1088/1741-2552/acd3b1.

18. Herff, Christian, Lorenz Diener, Miguel Angrick, Emily Mugler, Matthew C. Tate, Matthew A. Goldrick, Dean J. Krusienski, Marc W. Slutzky, and Tanja Schultz. 2019. “Generating Natural, Intelligible Speech From Brain Activity in Motor, Premotor, and Inferior Frontal Cortices.” Frontiers in Neuroscience 13 (November): 1267.

19. Herff, Christian, Dominic Heger, Adriana de Pesters, Dominic Telaar, Peter Brunner, Gerwin Schalk, and Tanja Schultz. 2015. “Brain-to-Text: Decoding Spoken Phrases from Phone Representations in the Brain.” Frontiers in Neuroscience 9 (June): 217.

20. Ingram, James N., Konrad P. Körding, Ian S. Howard, and Daniel M. Wolpert. 2008. “The Statistics of Natural Hand Movements.” Experimental Brain Research. Experimentelle Hirnforschung. Experimentation Cerebrale 188 (2): 223–36.

21. Jarosiewicz, Beata, Nicolas Y. Masse, Daniel Bacher, Sydney S. Cash, Emad Eskandar, Gerhard Friehs, John P. Donoghue, and Leigh R. Hochberg. 2013. “Advantages of Closed-Loop Calibration in Intracortical Brain–computer Interfaces for People with Tetraplegia.” Journal of Neural Engineering 10 (4): 046012.

22. Jarosiewicz, Beata, Anish A. Sarma, Daniel Bacher, Nicolas Y. Masse, John D. Simeral, Brittany Sorice, Erin M. Oakley, et al. 2015. “Virtual Typing by People with Tetraplegia Using a Self-Calibrating Intracortical Brain-Computer Interface.” Science Translational Medicine 7 (313): 313ra179.

23. Jorge, Ahmed, Dylan A. Royston, Elizabeth C. Tyler-Kabara, Michael L. Boninger, and Jennifer L. Collinger. 2020. “Classification of Individual Finger Movements Using Intracortical Recordings in Human Motor Cortex.” Neurosurgery 87 (4): 630–38.

24. Kao, Jonathan C., Paul Nuyujukian, Stephen I. Ryu, and Krishna V. Shenoy. 2017. “A High-Performance Neural Prosthesis Incorporating Discrete State Selection With Hidden Markov Models.” IEEE Transactions on Bio-Medical Engineering 64 (4): 935–45.

25. Koyama, Shinsuke, Steven M. Chase, Andrew S. Whitford, Meel Velliste, Andrew B. Schwartz, and Robert E. Kass. 2010. “Comparison of Brain-Computer Interface Decoding Algorithms in Open-Loop and Closed-Loop Control.” Journal of Computational Neuroscience 29 (1-2): 73–87.

26. Metzger, Sean L., Kaylo T. Littlejohn, Alexander B. Silva, David A. Moses, Margaret P. Seaton, Ran Wang, Maximilian E. Dougherty, et al. 2023. “A High-Performance Neuroprosthesis for Speech Decoding and Avatar Control.” Nature 620 (7976): 1037–46.

27. Microsystems, Blackrock. 2023. “NeuroPort Biopotential Signal Processing System: Instructions for Use.”

28. Moses, David A., Sean L. Metzger, Jessie R. Liu, Gopala K. Anumanchipalli, Joseph G. Makin, Pengfei F. Sun, Josh Chartier, et al. 2021. “Neuroprosthesis for Decoding Speech in a Paralyzed Person with Anarthria.” The New England Journal of Medicine 385 (3): 217–27.

29. Mugler, Emily M., James L. Patton, Robert D. Flint, Zachary A. Wright, Stephan U. Schuele, Joshua Rosenow, Jerry J. Shih, Dean J. Krusienski, and Marc W. Slutzky. 2014. “Direct Classification of All American English Phonemes Using Signals from Functional Speech Motor Cortex.” Journal of Neural Engineering 11 (3): 035015.

30. Nason, Samuel R., Matthew J. Mender, Alex K. Vaskov, Matthew S. Willsey, Nishant Ganesh Kumar, Theodore A. Kung, Parag G. Patil, and Cynthia A. Chestek. 2021. “Real-Time Linear Prediction of Simultaneous and Independent Movements of Two Finger Groups Using an Intracortical Brain-Machine Interface.” Neuron 109 (19): 3164–77.e8.

31. Nuyujukian, Paul, Jose Albites Sanabria, Jad Saab, Chethan Pandarinath, Beata Jarosiewicz, Christine H. Blabe, Brian Franco, et al. 2018. “Cortical Control of a Tablet Computer by People with Paralysis.” PloS One 13 (11): e0204566.

32. Pandarinath, Chethan, Paul Nuyujukian, Christine H. Blabe, Brittany L. Sorice, Jad Saab, Francis R. Willett, Leigh R. Hochberg, Krishna V. Shenoy, and Jaimie M. Henderson. 2017. “High Performance Communication by People with Paralysis Using an Intracortical Brain-Computer Interface.” eLife 6 (February). 10.7554/eLife.18554.

33. Pedregosa, Fabian, Gaël Varoquaux, Alexandre Gramfort, Vincent Michel, Bertrand Thirion, Olivier Grisel, Mathieu Blondel, et al. 2011. “Scikit-Learn: Machine Learning in Python.” Journal of Machine Learning Research: JMLR 12 (85): 2825–30.

34. Pitt, Kevin M., and Jonathan S. Brumberg. 2018. “Guidelines for Feature Matching Assessment of Brain-Computer Interfaces for Augmentative and Alternative Communication.” American Journal of Speech-Language Pathology / American Speech-Language-Hearing Association 27 (3): 950–64.

35. Povey, D., A. Ghoshal, and G. Boulianne. 2011. “The Kaldi Speech Recognition Toolkit.” Speech Recognition *…*. https://infoscience.epfl.ch/record/192584.

36. Scherer, Marcia J., Caren Sax, Alan Vanbiervliet, Laura A. Cushman, and John V. Scherer. 2005. “Predictors of Assistive Technology Use: The Importance of Personal and Psychosocial Factors.” Disability and Rehabilitation 27 (21): 1321–31.

37. Shah, Nishal P., Donald Avansino, Foram Kamdar, Claire Nicolas, Anastasia Kapitonava, Carlos Vargas-Irwin, Leigh R. Hochberg, et al. 2023. “Pseudo-Linear Summation Explains Neural Geometry of Multi-Finger Movements in Human Premotor Cortex.” bioRxiv. 10.1101/2023.10.11.561982.

38. Temmar, Hisham, Matthew S. Willsey, Joseph T. Costello, Matthew J. Mender, Luis H. Cubillos, Jordan Lw Lam, Dylan M. Wallace, Madison M. Kelberman, Parag G. Patil, and Cynthia A. Chestek. 2024. “Artificial Neural Network for Brain-Machine Interface Consistently Produces More Naturalistic Finger Movements than Linear Methods.” bioRxiv : The Preprint Server for Biology, March. 10.1101/2024.03.01.583000.

39. “The CMU Pronouncing Dictionary.” n.d. Accessed November 25, 2023. http://www.speech.cs.cmu.edu/cgi-bin/cmudict.

40. Vargas-Irwin, C. E., T. Hosman, J. Gusman, T K Pun, T Singer-Clark, A. Kapitonava, N P Shah, F. Kamdar, and L R Hochberg. 2022. “Single Hemisphere Encoding of 48 Right and Left Hand Gestures in Human Precentral Gyrus.” In 2022 Neuroscience Meeting Planner. Society for Neuroscience.

41. Willett, Francis R., Donald T. Avansino, Leigh R. Hochberg, Jaimie M. Henderson, and Krishna V. Shenoy. 2021. “High-Performance Brain-to-Text Communication via Handwriting.” Nature 593 (7858): 249–54.

42. Willett, Francis R., Darrel R. Deo, Donald T. Avansino, Paymon Rezaii, Leigh R. Hochberg, Jaimie M. Henderson, and Krishna V. Shenoy. 2020. “Hand Knob Area of Premotor Cortex Represents the Whole Body in a Compositional Way.” Cell 181 (2): 396–409.e26.

43. Willett, Francis R., Erin M. Kunz, Chaofei Fan, Donald T. Avansino, Guy H. Wilson, Eun Young Choi, Foram Kamdar, et al. 2023. “A High-Performance Speech Neuroprosthesis.” Nature 620 (7976): 1031–36.

44. Willsey, Matthew S., Samuel R. Nason-Tomaszewski, Scott R. Ensel, Hisham Temmar, Matthew J. Mender, Joseph T. Costello, Parag G. Patil, and Cynthia A. Chestek. 2022. “Real-Time Brain-Machine Interface in Non-Human Primates Achieves High-Velocity Prosthetic Finger Movements Using a Shallow Feedforward Neural Network Decoder.” Nature Communications 13 (1): 6899.

45. Willsey, Matthew S., Nishal P. Shah, Donald T. Avansino, Nick V. Hahn, Ryan M. Jamiolkowski, Foram B. Kamdar, Leigh R. Hochberg, Francis R. Willett, and Jaimie M. Henderson. 2024. “A Real-Time, High-Performance Brain-Computer Interface for Finger Decoding and Quadcopter Control.” bioRxiv. 10.1101/2024.02.06.578107.

46. Wodlinger, B., J. E. Downey, E. C. Tyler-Kabara, A. B. Schwartz, M. L. Boninger, and J. L. Collinger. 2015. “Ten-Dimensional Anthropomorphic Arm Control in a Human Brain-Machine Interface: Difficulties, Solutions, and Limitations.” Journal of Neural Engineering 12 (1): 016011.

47. Xu, Jing, Firas Mawase, and Marc H. Schieber. 2024. “Evolution, Biomechanics, and Neurobiology Converge to Explain Selective Finger Motor Control.” *Physiological Reviews*, February. 10.1152/physrev.00030.2023.

48. Young, D., F. Willett, W. D. Memberg, B. Murphy, B. Walter, J. Sweet, J. Miller, L. R. Hochberg, R. F. Kirsch, and A. B. Ajiboye. 2018. “Signal Processing Methods for Reducing Artifacts in Microelectrode Brain Recordings Caused by Functional Electrical Stimulation.” Journal of Neural Engineering 15 (2): 026014.

49. Yousry, T. A., U. D. Schmid, H. Alkadhi, D. Schmidt, A. Peraud, A. Buettner, and P. Winkler. 1997. “Localization of the Motor Hand Area to a Knob on the Precentral Gyrus. A New Landmark.” Brain: A Journal of Neurology 120 (Pt 1) (January): 141–57.

